# Ectopic gene conversion causing quantitative trait variation

**DOI:** 10.1101/2024.12.26.630369

**Authors:** Marina Pfalz, Seïf-Eddine Naadja, Jacqui Anne Shykoff, Juergen Kroymann

**Affiliations:** Ecologie Société Evolution, CNRS/Université Paris-Saclay/AgroParisTech, Gif-sur-Yvette, France

## Abstract

Why is there so much non-neutral genetic variation segregating in natural populations? We dissect function and evolution of a near-cryptic quantitative trait locus (QTL) for defense metabolites in Arabidopsis using the CRISPR/Cas9 system and nucleotide polymorphism patterns. The QTL is explained by genetic variation in a family of four tightly linked indole-glucosinolate *O*-methyltransferase genes. Some of this variation appears to be maintained by balancing selection, some appears to be generated by non-reciprocal transfer of sequence, also known as ectopic gene conversion (EGC), between functionally diverged gene copies. Here we elucidate how EGC, as an inevitable consequence of gene duplication, could be a general mechanism for generating genetic variation for fitness traits.

## Introduction

Mutation-selection balance explains the occurrence of genetic variation in populations: mutations arise continuously, and natural selection removes or fixes them at varying rates. However, natural populations harbor too much non-neutral genetic variation for mutation-selection balance alone to explain (Charlesworth 2015). Indeed, additional processes like migration and drift introduce or maintain variation in natural populations, and some variation is maintained by balancing selection (Charlesworth 2015; Kroymann and Mitchell-Olds 2005). Here we propose an additional, general mechanism for generating non-neutral variation. We elucidate how ectopic gene conversion (EGC) generates fitness variation in populations, illustrated by dissection of a quantitative trait locus for glucosinolates in Arabidopsis.

Gene duplication generates evolutionary novelty via specialization of different copies (Ohno 1970; Ohta 2000), optimizing related functions within a complex fitness landscape of similar yet distinct tasks (Force et al. 1999; Lynch and Conery 2000; Lynch and Force 2000; He and Zhang 2005). However, because these copies retain sequence similarity despite specialization, they are prone to unequal crossover and ectopic gene conversion (EGC). Unequal crossover will lead to variation in copy number and EGC, the non-reciprocal transfer of sequence between paralogous genes in the context of DNA repair, will homogenize variation among copies. Initially, in young gene copies, EGC is likely but will be largely neutral. As gene copies diverge in sequence and function, EGC will have a greater impact on phenotype but becomes less likely. Thus, it is easy to conclude that EGC can play only a very minor role in the evolution of gene families (Nei and Rooney 2006). Nonetheless, EGC has been documented across a wide range of organisms, from bacteria to humans (Ohta 1982; Innan 2003; Kroymann et al. 2003; Morris and Drouin 2004; Osada and Innan 2008; Benovoy and Drouin 2009; Arguello and Connallon 2011; Hanikenne et al. 2013; Harpak et al. 2017; Lamping et al. 2017) and is often cited as a mechanism for concerted evolution (Zimmer et al. 1980; Ohta 1991; Mano and Innan 2008).

We contend that EGC can also generate fitness variation that will be targeted by selection. When EGC places a specialized allele adapted to a particular cellular role into another, inappropriate context, this should produce maladapted variants. Here we describe this process, detected while dissecting a QTL for glucosinolate metabolism in Arabidopsis. Sequence alignments revealed clear traces of EGC among four tandemly arranged members of a gene family for defense metabolites. Using CRISPR technology to knock out different copies of the gene family we showed which allele in which context was responsible for phenotypic variation, confirming that EGC was the mechanism that generated functional variation.

Glucosinolates are amino-acid derived defense metabolites found in Brassicales. Due to their extensive intra- and interspecific genetic variation, they have become model compounds for understanding the genomic architecture of adaptive quantitative traits (Kliebenstein et al. 2001; Kliebenstein et al. 2005; Chan et al. 2011). Indole glucosinolates (IGs) are derived from tryptophan, are inducible by microbial pathogens or phytophagous insects, and play an important role in plant defense (Kim and Jander 2007; Bednarek et al. 2009; Clay et al. 2009; Pfalz et al. 2009; Pfalz et al. 2016). We had previously mapped leaf QTL for these IGs in Arabidopsis and cloned the *Indole Glucosinolate Modifier 1* (*IGM1*) QTL on chromosome 5 (Pfalz et al. 2007; Pfalz et al. 2009), which controls variation for 4-hydroxy-indol-3ylmethyl glucosinolate (4OHI3M) and 4-methoxy-indol-3ylmethyl glucosinolate (4MOI3M). A second QTL on chromosome 1, termed *Indole Glucosinolate Modifier 2* (*IGM2*), which we dissect here, was thought to control variation for 4OHI3M but not 4MOI3M (Pfalz et al. 2007).

A tightly linked gene family of indole glucosinolate *O*-methyltransferase genes (*IGMT*s) underlies *IGM2*, via a complex determination of metabolite levels. The gene family presents multiple cases of shared polymorphisms that are unlikely to have arisen by independent point mutations. Instead, we contend that these occurred through EGC, and that the counterplay between selection and EGC shaped the QTL.

## Results

### Mapping and dissection of *IGM2*

We fine-mapped *IGM2* using F_2_ progeny from a cross between two near-isogenic lines (NILs) of the Arabidopsis Da(1)-12 × Ei-2 mapping population, DE096 and DE155 (Pfalz et al. 2007). In leaves, DE096 had more 4OHI3M than DE155, but the NILs did not differ in their 4MOI3M content. In roots, DE096 had more 4OHI3M and less 4MOI3M than DE155 (**Fig. S1a**). DE096 and DE155 differed only in a ∼ 20 Mbp region containing *IGM2*, while their marker genotypes were identical for the rest of the genome (**Fig. S1b**). We phenotyped for the fraction of 4OHI3M in the combined amount of 4OHI3M and 4MOI3M. *IGM2* mapped to a ca. 1.6 Mbp interval, with no evidence of additional QTL nearby (**Figs. S1c and S1d**). Statistical support for a QTL centered near a family of four tandemly arranged methyltransferase genes. These genes were previously identified as coding for indole glucosinolate *O*-methyltransferases (IGMTs) that convert 4OHI3M to 4MOI3M (Pfalz et al. 2011), making them plausible candidates for the QTL (**Figs. S1e and S1f**).

We conducted quantitative real-time PCR to assess whether differences in *IGMT* expression could explain the QTL. We used primers specific to the target genes, ensuring substantial mismatches with other gene family members while avoiding polymorphisms at primer binding sites between accessions (**Table S1**). Additionally, we employed a primer pair that could simultaneously amplify all genes. Leaf *IGMT1* and *3* transcript levels were higher in DE155 than in DE096, as was the overall transcript level, while *IGMT2* and *4* transcript levels did not differ between the two NILs (**Fig. 1**). In roots, *IGMT1* transcript levels were higher in DE155, but *IGMT3*, *4*, and particularly total *IGMT* transcripts were lower. Thus, genotypic differences in *IGMT* transcript abundance alone could not explain the observed quantitative metabolite pattern.

**Fig. 1.**
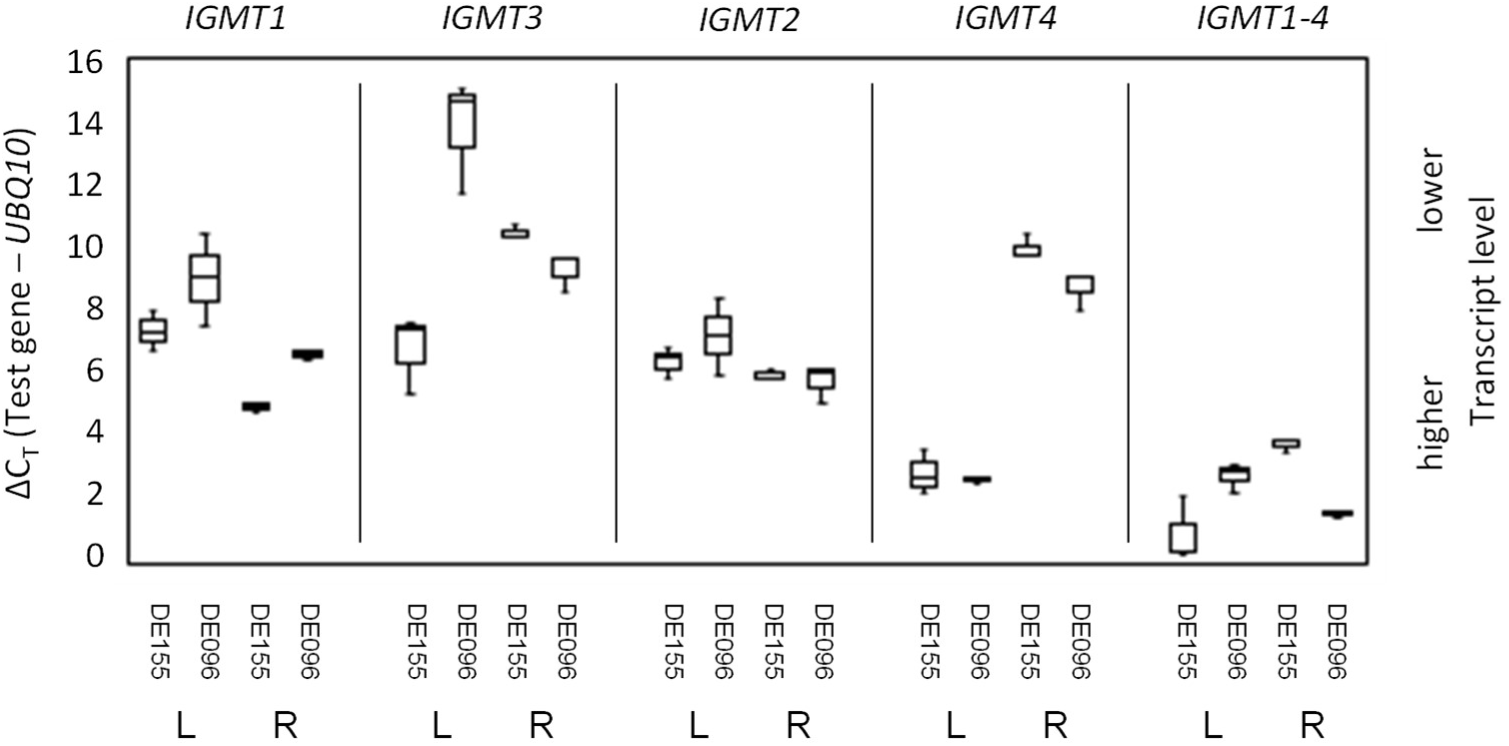
*IGMT* transcript levels in DE155 and DE096. Shown are boxplots with maximum, median and minimum of ΔC_T_ values for leaves (L) and roots (R) in comparison to a control gene, *UBQ10*. Each genotype/transcript combination had n = 3 biological samples. *IGMT1* to *IGMT*4 represent individual genes, while *IGMT1-4* indicates total *IGMT* transcript levels. Note that higher ΔC_T_ values correspond to lower expression levels, and *vice versa*.

Therefore, we employed CRISPR/Cas9 to generate *igmt* mutants in both NILs. Our approach involved a construct that produced three single guide RNAs (sgRNAs), each targeting distinct sites within the four *IGMT* sequences. However, due to sequence variation, the sgRNAs did not match precisely in every instance. We Sanger-sequenced all Cas9 target sites within the four *IGMT* genes in our mutant lines. Typically, we detected small insertions or deletions that caused frameshifts. However, in certain cases, we observed larger deletions resulting in the fusion of mismatched sequence from neighboring genes (**Fig. S2**).

We obtained quadruple knockouts, *igmt1-4*_DE155_ and *igmt1-4*_DE096_, for both NILs. While the leaves of these mutants lacked 4MOI3M entirely, the roots still exhibited a background level of 4MOI3M, suggesting the presence of another enzyme capable of converting 4OHI3M to 4MOI3M in roots. Our prime candidate was IGMT5, encoded on chromosome 5, which normally converts 1-hydroxy-indol-3ylmethyl to 1-methoxy-indole-3ylmethyl glucosinolate and has high root but low leaf activity (Pfalz et al. 2016). We therefore generated an *igmt1-4* quadruple knockout in the *igmt5* mutant background. As expected, these plants showed no detectable 4MOI3M in either leaves or roots (**Table 1**).

**Table 1.** Estimated Marginal Means - 4MOI3M/(4OHI3M + 4MOI3M) in Roots.

| Genotype | Mean | SEM | 95% Confidence Interval |  |
| --- | --- | --- | --- | --- |
|  |  |  | Lower | Upper |
| DE155 | 0.533 | 0.019 | 0.495 | 0.570 |
| *igmt1-4*<sub>DE155</sub> | 0.050 | 0.015 | 0.020 | 0.080 |
| DE096 | 0.361 | 0.017 | 0.326 | 0.396 |
| *igmt1-4*<sub>DE096</sub> | 0.035 | 0.014 | 0.006 | 0.063 |
| Col-0 | 0.297 | 0.017 | 0.264 | 0.330 |
| *igmt5*<sub>Col-0</sub> | 0.260 | 0.019 | 0.223 | 0.297 |
| *igmt1-5*<sub>Col-0</sub> | 0.000 | 0.015 | -0.030 | 0.030 |

8 ≤ n ≤ 12

In addition to the quadruple mutant, referred to as *igmt1-4*_DE155_, we obtained the genotypes *igmt3*_DE155_, *igmt1/3*_DE155_, and *igmt1-3*_DE155_ for DE155, and *igmt1/4*_DE096_, *igmt1/3/4*_DE096_, and *igmt1/2/4*_DE096_ for DE096 **(Fig. S2)**.

In leaves, we observed the most substantial differences in the levels of 4OHI3M, 4MOI3M and/or the fraction of 4OHI3M when comparing *igmt1-3*_DE155_ with *igmt1-4*_DE155_, and DE096 wildtype with *igmt1/4*_DE096_ (**Fig. 2**). Thus, *IGMT4* had the largest impact on 4MOI3M generation in the leaves of both NILs. Additionally, we found noticeable differences between *igmt1/2/4*_DE096_ and *igmt1/4*_DE096_, and between *igmt1/3/4*_DE096_ and *igmt1-4*_DE096_ (**Fig. 2**), which highlighted the role of *IGMT2*_DE096_. In stark contrast, we only found minor differences for *IGMT2*_DE155_ when comparing *igmt1/3*_DE155_ and *igmt1-3*_DE155_, indicating that genotypic variation in *IGMT2* contributed to the leaf QTL.

**Fig. 2.**
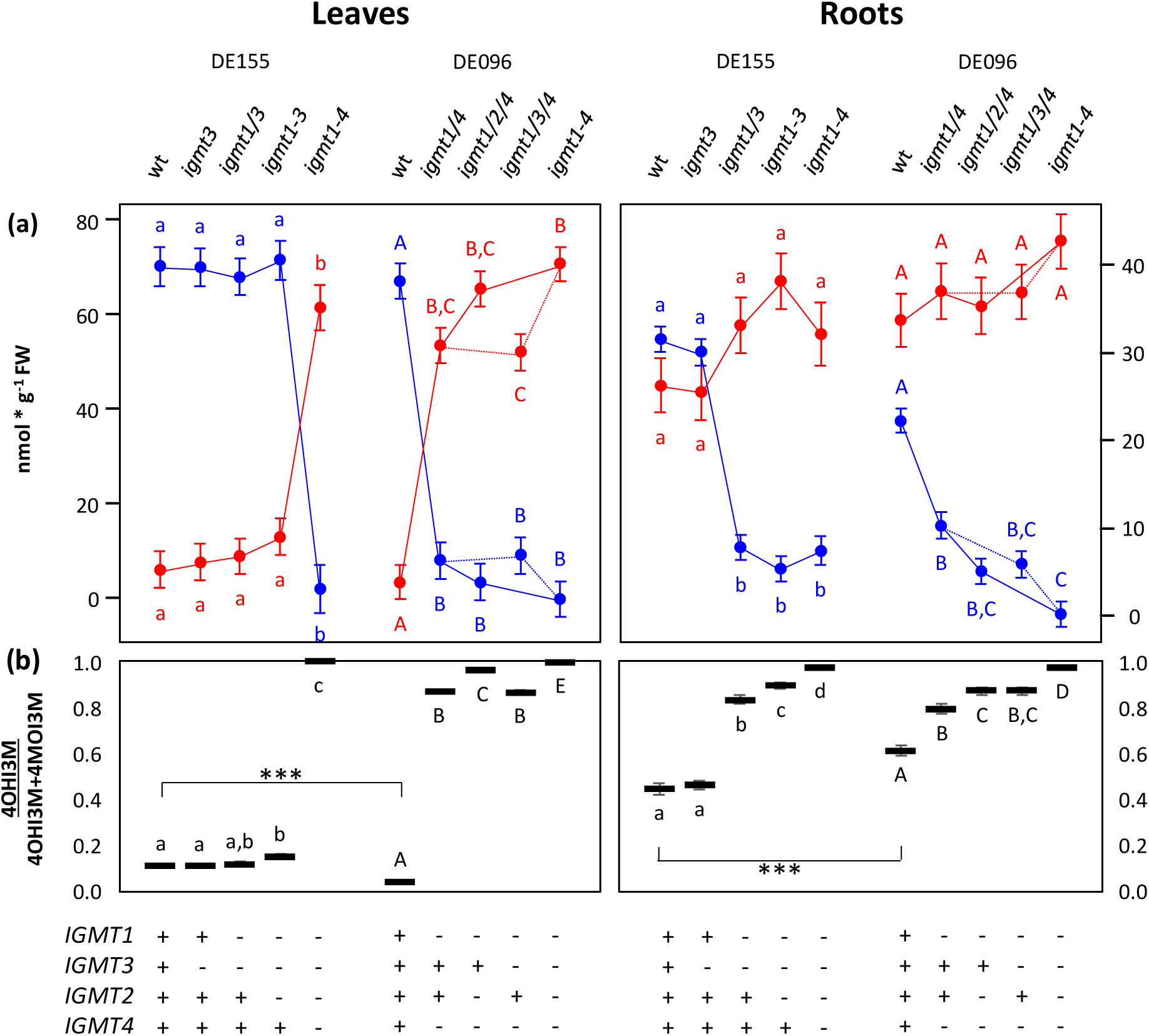
Quantitative differences in 4MOI3M and 4OHI3M in leaves and roots. Shown are estimated marginal means of metabolite concentration in nmol per gram fresh weight (± SEM) for Arabidopsis DE155, DE096 and *igmt* mutants. Different upper and lower case letters indicate statistically significant differences. Each genotype had n = 10 or 11 samples. a) Concentration of 4MOI3M (blue) and 4OHI3M (red). Lines connect meaningful comparisons. b) Retransformed data for the fraction of 4OHI3M in the combined amount of 4OHI3M and 4MOI3M. Genotypes at *IGMT 1 – 4* are indicated, with “-” for defective and “+” for functional genes. NILs differ significantly for the fraction of 4OHI3M, indicated by asterisks.

However, given that the generation of each 4MOI3M molecule consumes one 4OHI3M molecule, the contrast between an active IGMT2 in DE096 and a largely inactive IGMT2 in DE155 alone could not explain the leaf QTL pattern. We therefore inspected the effects of IGMT4 in both NILs more closely, based on the combined analysis of data for *igmt1-3*_DE155_, *igmt1-4*_DE155_, *igmt1/4*_DE096_, and DE096 from four independent experiments. Indeed, this analysis unveiled an additional quantitative difference between the NILs (**Fig. 3a**). While the increase in 4OHI3M was consistent in the comparisons from *igmt1-3*_DE155_ to *igmt1-4*_DE155_ and from DE096 to *igmt1/4*_DE096_, the reduction in 4MOI3M was significantly more pronounced when *IGMT4* was knocked out in the DE155 background. This indicated that IGMT4 had a higher level of activity in DE155 compared to DE096, thereby increasing the flux through the pathway.

**Fig. 3.**
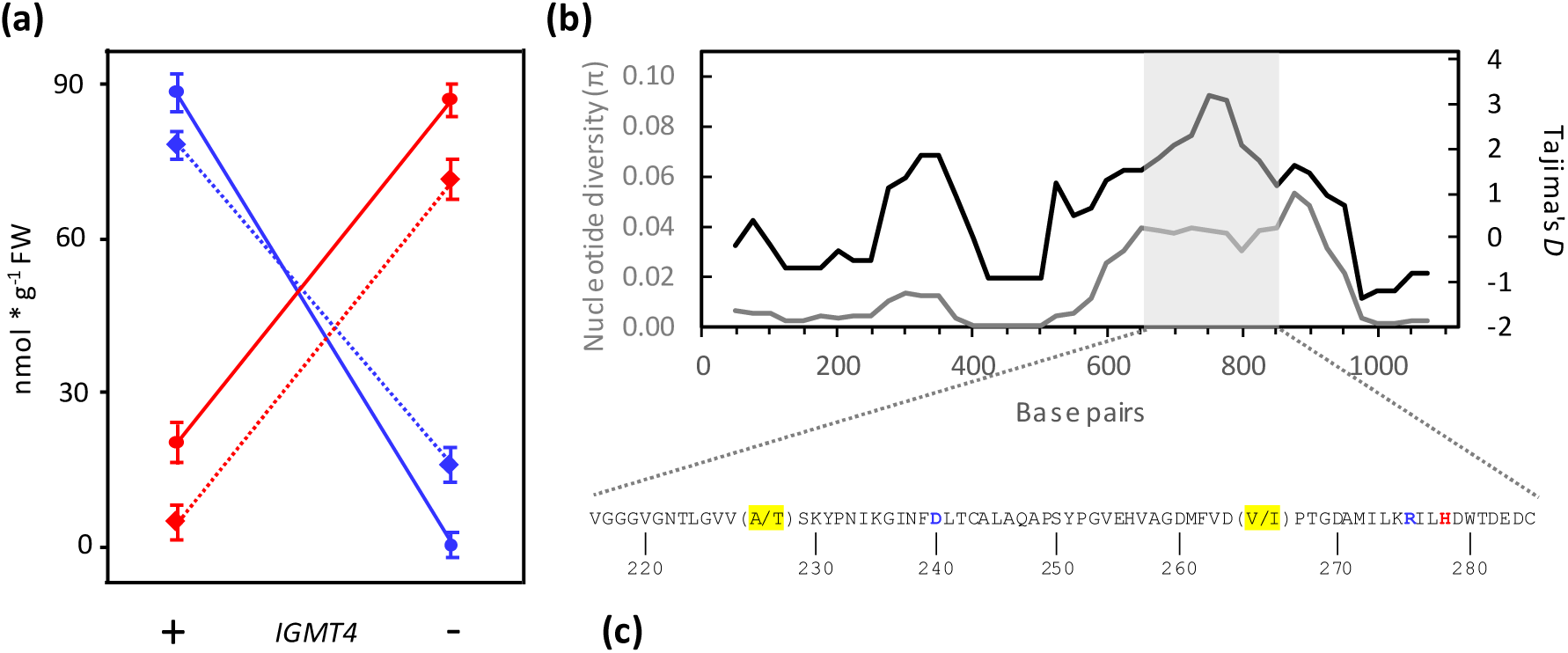
Analysis of *IGMT4*. a) Quantitative effects of *IGMT4* on leaf IGs in DE155 and DE096. Data pooled from four independent experiments show estimated marginal means of metabolite concentration in nmol per gram fresh weight (± SEM) of 4OHI3M (red) and 4MOI3M (blue) with *IGMT4* intact (left) or defective (right) in DE155 (circles and solid lines) and DE096 (diamonds and dashed lines). Each genotype had n = 35 – 61 samples. The interaction of *IGMT4* by background is highly significant for 4MOI3M (*F*_1,164_ = 18.7; *p* < 0.0001) but not for 4OHI3M (*F*_1,164_ = 0.0; *p* = 0.994). There were no significant genotype or genotype-by-background effects for I3M, 1MOI3M, and total IG. b) Sliding window analyses of nucleotide polymorphisms (gray) and Tajima’s *D* (black) with a window size of 100 and a step size of 25 nucleotides. Tajima’s *D* is significantly elevated (*D* > 2.07; *p* < 0.05) from ca. 650 to ca. 850 bp (shaded gray). c) Functionally important amino acids. S-adenosyl-L-methionine binding sites are shown in blue, and the active site in red. Two polymorphic amino acids are highlighted yellow.

We also conducted tests to ascertain if there were differences in the levels of *CYP81F* transcripts between DE155 and DE096, or if mutations in *IGMT* genes influenced the transcript abundance of wildtype *IGMT*s. However, we observed no substantial alterations (**Fig. S3**).

Based on these results, we could explain the leaf QTL as the cumulative effects of two distinct quantitative trait genes (QTGs), namely *IGMT2* and *IGMT4*. Genotypic variation in *IGMT4* led to an excess of 4MOI3M in DE155 without altering 4OHI3M levels. This should result in a QTL for 4MOI3M, with DE155 having the high allele for *IGMT4*. However, genotypic variation in *IGMT2* resulted in an elevation of 4MOI3M in DE096, which counterbalanced the excess of 4MOI3M in DE155, and concurrently caused a reduction in 4OHI3M. Consequently, the collective impact of genotypic variation in both QTGs manifested as a disparity in 4OHI3M levels, with no noticeable difference in the levels of 4MOI3M.

In the roots of DE155, *IGMT1* was the primary contributor to 4MOI3M biosynthesis, as evidenced by the contrast between *igmt3*_DE155_ and *igmt1/3*_DE155_ (**Fig. 2**). *IGMT4* and, notably, *IGMT2* also played roles, but *IGMT3* had no discernible impact. In the roots of DE096, we noted the largest contrast between *igmt1/4*_DE096_ and DE096 wildtype. However, the *igmt1/4*_DE096_ double mutant alone did not allow us to discern whether the observed effects were attributable to *IGMT1*_DE096_, *IGMT4*_DE096_, or their combination. Furthermore, *IGMT2*_DE096_ and *IGMT3*_DE096_ were implicated in the conversion of 4OHI3M to 4MOI3M, primarily observed through the impact of the corresponding mutants on the fraction of 4OHI3M.

The influence of *IGMT1*_DE155_ was substantially more pronounced than the combined effect of *IGMT1*_DE096_ and *IGMT4*_DE096_. Hence, *IGMT1* was a root QTG. *IGMT3* was a second, but cryptic, root QTG, with DE096 possessing the high allele that mitigated the impact of *IGMT1*_DE155_. In contrast to leaves, genotypic variation in *IGMT2* had no noticeable quantitative effect on roots.

This root QTL pattern corresponded well with the abundance of *IGMT* transcripts in both NILs (**Fig. 1**), suggesting that quantitative variation in the roots was mainly governed by differences in the expression of individual genes. The transcript level of *IGMT1* was higher and that of *IGMT3* lower in DE155 than in DE096, while the transcript level of *IGMT2* was similar between NILs. Interestingly, *IGMT4*_DE096_ exhibited a higher transcript abundance than *IGMT4*_DE155_, indicating that *IGMT4* could be another cryptic root QTG, further counteracting the effect of genotypic variation in *IGMT1*.

### Patterns of variation in the *IGMT* cluster

In the comprehensive assessment of QTL effects and transcript level variation among NILs, *IGMT2*_DE155_ stood out. The DE155 allele was obviously functional in roots, contributing to 4MOI3M at a level comparable to its DE096 counterpart (**Fig. 2**). However, *IGMT2*_DE155_ activity in leaves was relatively minor, despite having a transcript level that was similar, if not slightly higher, than that of *IGMT2*_DE096_ (**Fig. 1**).

To investigate the cause of this peculiar phenomenon, we built a gene tree with all four genes from both NILs. We expected alleles of the same gene copies to cluster together, which was indeed the case for *IGMT1* and *IGMT4*. Much to our surprise, however, the relationship between *IGMT2* and *IGMT3* genes and alleles remained unresolved (**Fig. S4**), suggesting that *IGMT2*_DE155_ and *IGMT3*_DE155_ did not evolve independently.

To examine this further, we procured and manually aligned sequences of the *IGMT* region from the 1001 Arabidopsis Genomes project (1001genomes.org), concentrating on *de novo*, reference-quality assembles (Jiao and Schneeberger 2020), and incorporated additional short-read assemblies with near-complete *IGMT* sequences (Gan et al. 2011) (**Table S2**). Our alignment contained data from 28 accessions, including several relict lineages (Toledo et al. 2020). All assemblies of the *IGMT* region, particularly those from long reads, showed exactly four *IGMT* copies, suggesting that this configuration exists since more than 200,000 generations (Durvasula et al. 2017).

We compared the coding sequences of all four *IGMT* genes and found numerous polymorphisms (**10.6084/m9.figshare.28093955**). To ensure these were not sequencing errors, we amplified and sequenced *IGMT4* coding sequence from 16 Arabidopsis accessions, confirming our findings. Strikingly, many of these polymorphisms were not copy-specific, but shared by two or even more gene copies, suggesting EGC (Mansai and Innan 2010).

We tested whether point mutations alone could explain the high number of cases of ‘shared polymorphisms’, where the same polymorphism segregated at the same site in two or more copies. We focused on instances where two independent point mutations affected the same site, either within a single gene copy or across two distinct copies. The relative probabilities of these events depended on the number of gene copies (n), being (n^-1^) for two mutations at the same site within a single gene copy and (1 - n^-1^) for two mutations at the same site across two distinct copies.

We assumed a large population with initially no polymorphisms within copies but allowing for fixed differences between gene copies. When the same gene copy mutates twice at the same site, two outcomes are possible: either both mutations lead to the same polymorphism, appearing as a single, copy-specific polymorphism, or they result in three different nucleotides segregating at the same site. When two mutations occur at the same site in two different copies, this can lead to a shared polymorphism or to two different copy-specific polymorphisms. The probabilities of these outcomes depend on whether the copies initially had identical or different nucleotides at the side in question, and in the latter case, whether the nucleotides differed by a transition or a transversion (see **Supplementary text** for details). From our data, we estimated a transition/transversion ratio of 2.5935, consistent with other studies (Ossowski et al. 2010; Weng et al. 2019). However, we were uncertain about the initial count of identical *versus* different positions in gene copy comparisons. Therefore, we examined the entire frequency spectrum of identical *versus* different positions, ranging from zero to 100%, considering their mutual exclusivity.

For n = 4 copies and a transition/transversion rate of 2.5935, as in the case of the Arabidopsis *IGMT* genes, the proportion of shared among visible types of polymorphisms can never exceed 50% (**Fig. S5**). At higher gene copy numbers, this proportion plateaus at 56%. In stark contrast, our entire dataset had only three cases with three different segregating nucleotides in the same gene copy and ten cases with two different segregating polymorphisms across multiple gene copies. However, shared polymorphisms across copies occurred at 61 sites, making up over 80% of the total visible polymorphisms that affected either the same copy twice or two different copies once each.

Thus, observed polymorphism patterns revealed far too many shared polymorphisms across gene copies than would be expected from independent point mutations, making EGC the most likely explanation. Indeed, visual inspection identified numerous linked specific variants at polymorphic sites, sometimes spanning hundreds of positions, indicative of gene conversion tracts. These included untranslated regions, coding sequences, and introns (**Tables S3** and **S4**). Reexamining the 13 cases of three segregating nucleotides in the same copy or with two different segregating polymorphisms across multiple gene copies revealed that most of these events were better explained by EGC than by independent point mutations. Hence, shared polymorphisms resulting from two independent point mutations were even more rare than initially suspected. Thus, shared polymorphisms overall indicate EGC.

*IGMT4*, the QTG where the knockouts varied for 4MOI3M but not for 4OHI3M, displayed high levels of intermediate frequency polymorphisms within a region of about 200 bp (**Fig. 3b**). This sequence stretch encompassed codons encoding functionally important amino acids, including S-adenosyl-L-methionine binding sites and the active site, corresponding to amino acids D240, R275 and H278, respectively (**Fig. 3c**). A sliding window analysis, using a window size of 100 and a step size of 25, found significantly elevated Tajima’s *D* (Tajima 1989) in this region of *IGMT4* but not in the other three gene copies. Similarly, the HKA test (Hudson et al. 1987) revealed a significant excess of intermediate frequency polymorphisms in the C-terminal half of *IGMT4* (*χ²* = 6.16, *p* < 0.05) compared to *IGMT*s from *A. lyrata*. The results of both tests support the presence of balancing selection.

### EGC and subfunctionalization

What are the consequences of EGC for the *IGMT* gene cluster? All four gene products performed the same reaction – converting 4OHI3M to 4MOI3M – but showed organ-specific differences in transcript abundance (**Fig. 1**). CRISPR/Cas9-induced mutations in one gene were not effectively offset by the remaining gene copies, which implied that different *IGMT*s may be expressed in distinct cells. Indeed, *IGMT2* and *IGMT3* are expressed in different root tissues (Cao et al. 2024). Together, these findings suggest subfunctionalization, a process that preserves gene duplicates in a genome by partitioning the functions of the ancestral gene (Force et al. 1999; Lynch and Conery 2000; Lynch and Force 2000; He and Zhang 2005). Subfunctionalization allows different gene copies to express in different contexts and to acquire mutations that optimize their function within these contexts, such as specific cellular environments. However, when EGC transfers these mutations to a gene copy specialized for a different cellular environment, they may impair the function of this copy, as exemplified by *IGMT2*_DE155_ in leaves (**Fig. 2**). Consequently, amino acids that enhance the environment-specific function of gene copies should be subject to purifying selection.

To test this, we focused on fixed derived amino acids identified by comparing *A. thaliana* and *Arabidopsis lyrata* (Hu et al. 2011) IGMT sequences. We divided each *A. thaliana IGMT* coding sequence into intervals delimited by adjacent polymorphisms shared with any of the other three gene copies. We then compared log_10_-transformed interval lengths with and without codons specifying fixed derived amino acids, using two-sided *t* tests. Intervals with one or more fixed derived amino acids were significantly larger than those without for *IGMT1* (*t_df_*_=19_ = −3.12, *p* < 0.01), *IGMT3* (*t_df_*_=46_ = −2.77, *p* < 0.01), *IGMT2* (*t_df_*_=42_ = −2.88, *p* < 0.01), and *IGMT4* (*t*_df=28_ = −2.42, *p* < 0.05) (**Fig. 4 and Fig. S6**). Across all four genes, results were highly significant (*t_df_*_=141_ = −5.72, *p* < 0.0001), suggesting that EGC was indeed selected against around those amino acids.

**Fig. 4.**
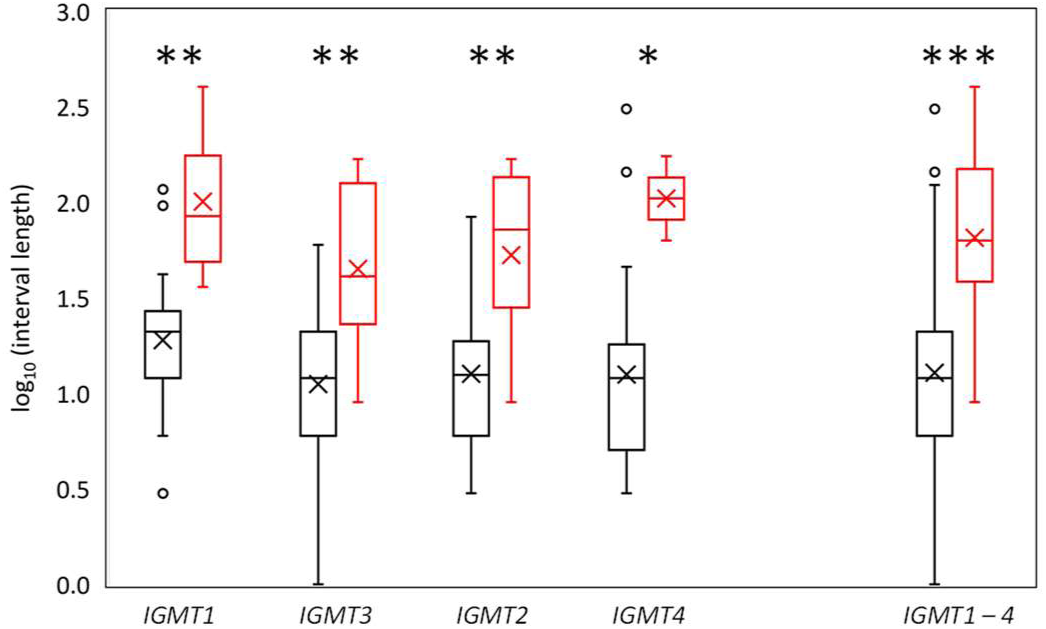
Boxplot comparing the log10-transformed interval sizes without (black) and with (red) fixed derived amino acids. These derived amino acids are nonsynonymous substitutions that distinguish Arabidopsis IGMTs from those in *Arabidopsis lyrata*. Each interval is flanked by neighboring shared polymorphisms, defined as the same nucleotides segregating at the same position in two or more gene copies. Shown are mean (crosses), median (black horizontal line), and range of the data including outliers (open circles). Statistical significance is indicated by asterisks (*: p < 0.05; **: p < 0.01; ***: p < 0.001).

## Discussion

We used the CRISPR/Cas9 system to dissect a near-cryptic QTL for defense metabolites. We obtained a set of different combinations of *IGMT* mutants in three genetic backgrounds but we did not obtain all possible knockout combinations. Consequently, we cannot entirely rule out that some observed effects result from interactions between gene products, such as heterodimers. However, the tissue-specific expression of these genes, coupled with the lack of coordinated expression and compensation for defective genes at the transcript level, argue against this possibility. Our analyses show that *IGM2* comprises a cluster of QTGs, with DE155 and DE096 both having high and low alleles, but at different gene copies. Thus, even small-effect QTL can have a complex genetic architecture.

To demonstrate the existence of EGC among *IGMT* copies, we counted sites with multiple polymorphisms, either within the same gene copy or across different copies. This approach provided a convenient means to assess EGC without requiring prior knowledge of mutation, recombination, or gene conversion rates. Independent point mutations are highly unlikely to generate polymorphisms shared across two or more gene copies, whereas EGC generates such shared polymorphisms, especially visible when conversion tracts encompass long sequence stretches with multiple nucleotide substitutions. Quantitative differences in IGMT performance were partially attributable to EGC, as shown by comparing IGMT2 between two near-isogenic Arabidopsis lines. Both alleles were equally active in roots, but IGMT2 from DE155, which had a clear signature of EGC, performed worse in leaves compared to its DE096 counterpart. Furthermore, EGC events have shaped polymorphism patterns across the entire *IGMT* gene cluster (**Fig. S7**), evidenced by sequence comparison of 28 Arabidopsis accessions, suggesting that other *IGMT* gene copies in other accessions may also have experienced performance alterations due to EGC.

Duplicated genes can persist longer than expected when higher gene dosage is advantageous (Sugino and Innan 2006; Hanikenne et al. 2013; Heidel-Fischer et al. 2019) or when they functionally diverge. Subfunctionalization is initially a neutral process during which gene duplicates partition the roles of the ancestral gene (Force et al. 1999, Stolzfus 1999). Consequently, if the presence of subfunctionalized copies is fixed by drift, selection preserves this configuration. Subfunctionalized copies may acquire advantageous mutations that enhance copy-specific function or confer novel function.

In all of these cases selection stabilizes the presence of multiple gene copies. For dosage effects, EGC leads to homogenization, and perhaps rapid spread of advantageous mutations between copies (Thomas 2006; Mano and Innan 2008; Hanikenne et al. 2013). Maladapted variants arise by mutation but are removed by selection. However, if gene copies have functionally diverged, EGC, as an unavoidable consequence of having multiple gene copies, introduces sequence from a copy adapted to a specific context into another copy adapted to a different context, as exemplified here. Thus, EGC generates inferior variants, which, within a population, manifest as QTL segregating fitness variation (**Fig. 5**). By stabilizing the presence of multiple copies and favoring their specialization, natural selection generates the opportunity for maladaptations to arise by EGC, until copies have sufficiently diverged to prevent EGC. Thus, natural selection itself sets the stage for the generation of more fitness variation than is explained by mutation-selection balance.

**Fig. 5.**
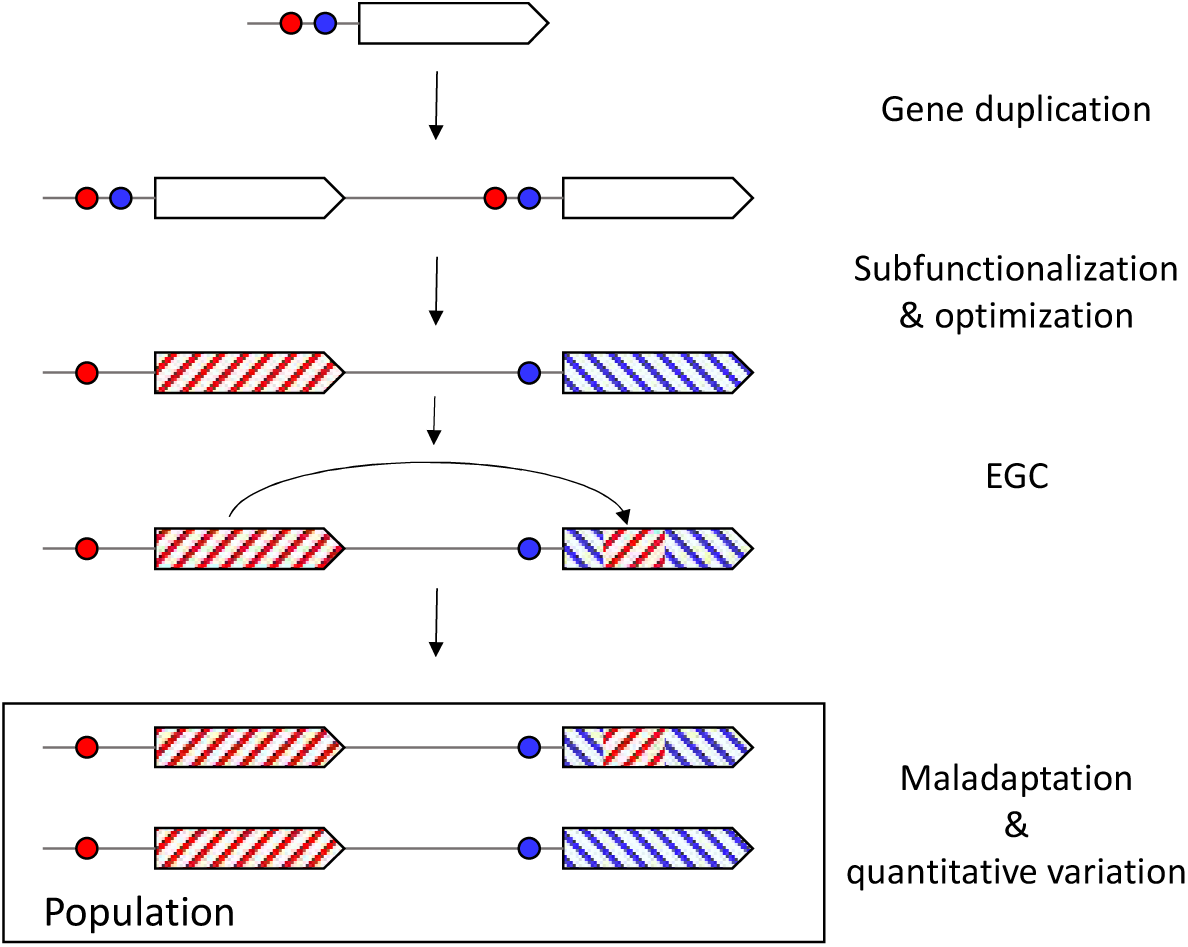
Maladaptation caused by EGC in gene families. After gene duplication, the paralogs acquire specific expression patterns and optimize accordingly. EGC introduces sequence from one copy optimized for one context into another copy optimized for a different context, leading to maladaptation. This results in quantitative genetic variation in the population.

## Materials and Methods

### Culture conditions

Arabidopsis plants were cultivated in growth chambers with 11.5 hours of light at 22**°**C and 12.5 hours of darkness at 16**°**C, maintaining approximately around 70 – 80% relative humidity. Seeds were sown on soil and stratified for three days at 6**°**C. For glucosinolate analysis, one week after germination, plants were transferred to sand and grown randomized in 96-celled trays, fertilized weekly with Hydrocani C2 liquid fertilizer (Hydro Agri), as described by (Pfalz et al. 2016).

### Glucosinolate extraction and analysis

Four weeks after germination, approximately 100 g leaf material and the entire root were harvested, weighed and snap-frozen in liquid nitrogen. Glucosinolate extracts were separated in their desulfo-form on a Vanquish HPLC system (Thermo), utilizing a LiChrospher 100 RP-18e LiChroCART column (250 x 4, 100A, 5 µm) as described by (Pfalz et al. 2011). Glucosinolate identification was based on retention time and UV spectra, and quantification was based on integrated absorption peak area at 229 nm, using sinigrin as an external standard and published response factors (Buchner 1987) to correct for different UV absorption capacities of various IGs.

### RNA extraction and quantitative real-time quantitative PCR (RT-qPCR)

Frozen leaf or root material was finely ground using a TissueLyser II (Qiagen). Total RNA was extracted with TriSure (Bioline) or the NucleoSpin RNA kit (Macherey & Nagel), treated with Turbo DNase (Ambion), and purified using RNeasy MinElute columns (Qiagen) or RNA Clean & Concentrator 5 (Zymo Research). RNA quality and quantity were assessed by agarose gel electrophoresis and a Nanodrop 2000 (Thermo). First strand cDNA synthesis was performed with the Maxima First Strand cDNA Synthesis Kit (Thermo), using 500 ng of total RNA. RT-qPCR was conducted with SYBR Green qPCR Master Mix (Thermo) on a StepOnePlus Real Time PCR System (Applied Biosystems), using Arabidopsis *UBQ10* (At4g05320) as the reference gene. Each genotype had three to six biological replicates. Primer sequences are listed in **Table S1**.

### Generation of mutant plants

Coding sequences of *IGMT1 – 4* from Arabidopsis accessions Col-0, Da(1)-12 and Ei-2 were used to design three single guide RNA (sgRNA) sequences targeting all four genes simultaneously. The design was performed using the CRISPOR software (crispor.tefor.net), with a preference for sites close to the 5’-end of the target genes and low off-target probability. A 1026 bp cassette flanked by attB sites, including all three sgRNAs, each driven by an *At*U6-26 promoter and containing the tracrRNA scaffold as well as the U6 terminator, was synthesized by Genewiz (Leipzig, Germany) and cloned into the pDONR207 vector (Thermo). The final cloning step into pDE-Cas9 DsRed (Fauser et al. 2014; Morineau et al. 2016), permitting selection by DsRed marker fluorescence, was achieved using Gateway^TM^ LR recombination. This vector was then used to transform *Agrobacterium tumefaciens* GV3101. The floral dip method (Clough and Bent 1998) was employed to generate mutants in Arabidopsis DE096, DE155, Col-0 and *igmt5*. After DNA extraction, mutations were identified by Sanger sequencing, using gene-specific primers (**Table S1**) covering the target regions of all three sgRNAs.

### Statistical and computational analyses

All measurements of were taken from distinct samples. Fine-mapping of *IGM2* was performed using a general linear model in Systat V9, with markers and growth trays as fixed factors. Glucosinolate quantities were analyzed via ANCOVA in jamovi Version 2.3.28.0 (www.jamovi.org), using the weight of the harvested plant tissue as a covariate and genotype as a fixed factor. When pooling data from multiple experiments, experiment was included as an additional fixed factor, along with an interaction term for genotype × experiment. The leaf effect of *IGMT4* was assessed by ANCOVA, with leaf weight as a covariate and genetic background, genotype at *IGMT4* and experiment as fixed factors. The data for the fraction of 4OHI3M in the combined amount of 4OHI3M and 4MOI3M were arcsine square root-transformed before ANOVA, with genotype as a fixed factor. Statistical differences between genotypes were assessed using post-hoc two-sided *t*-tests. PAML (Yang 2007) was used to estimate the transition/transversion ratio in *IGMT* genes. SplitsTree (Huson and Bryant 2006) was employed to construct a genealogical network for Arabidopsis *IGMT* sequences. DnaSP v6 (Rozas et al. 2017) was used for sliding window analyses of nucleotide polymorphisms and of Tajima’s *D* and for the HKA test. Functionally important sites in IGMT4 were identified using ScanProsite (De Castro et al. 2006) on the Expasy webserver (www.expasy.org).

## Acknowledgements

We are grateful for funding from the Agence Nationale de la Recherche (ANR-10-GENM-005, ANR-20-CE92-0042) (J.K.). We thank Marine Paupiére for her contribution to QTL mapping and George Sandler for helpful comments.

## Author contributions

M.P. and S.-e. N. performed experiments and analyzed data. J.K. conceived the study and analyzed data. J.K. and J.S. wrote the manuscript with contributions from M.P.

## Competing interests

Authors declare that they have no competing interests.

## Data and materials availability

All data are available in the main text or the supplementary materials.

## Supplementary Materials

Supplementary Text

Figs. S1 to S7

Tables S1 to S4

## Supplementary Materials

### Supplementary text

#### Calculations of probabilities for different types of nucleotide polymorphisms

We consider a large population with n copies of a gene. Initially, there is no polymorphism within copies but fixed differences among copies are allowed. We examine cases where two independent point mutations affect the same site, either twice in the same copy or once in two different copies. The probabilities of these events are P_same copy_ = n^-1^ and P_different copies_ = (1 – n^-1^), respectively.

(2 copies: AA*, AB, BA, BB*

3 copies: AA*, AB, AC, BA, BB*, BC, CA, CB, CC*

4 copies: AA*, AB, AC, AD, BA, BB*, BC, BD, CA, CB, CC*, CD, DA, DB, DC, DD*)

Each nucleotide can change into any of the three other nucleotides. There is one possibility for a transition, t_i_ (*e.g.*, A -> G) and two possibilities for a transversion, t_v_ (*e.g.*, A -> C, A -> T). The probability for a transition is P_ti_ = x, the probability for each transversion is ½ P_tv_ = ½ (1 – x). The transition/transversion ratio t_i_/t_v_ can be estimated from the data.

We first determine the probability that the same mutation occurs twice at the same site, either in one copy or in two different copies, with µ_1_ being the first and µ_2_ being the second mutation:

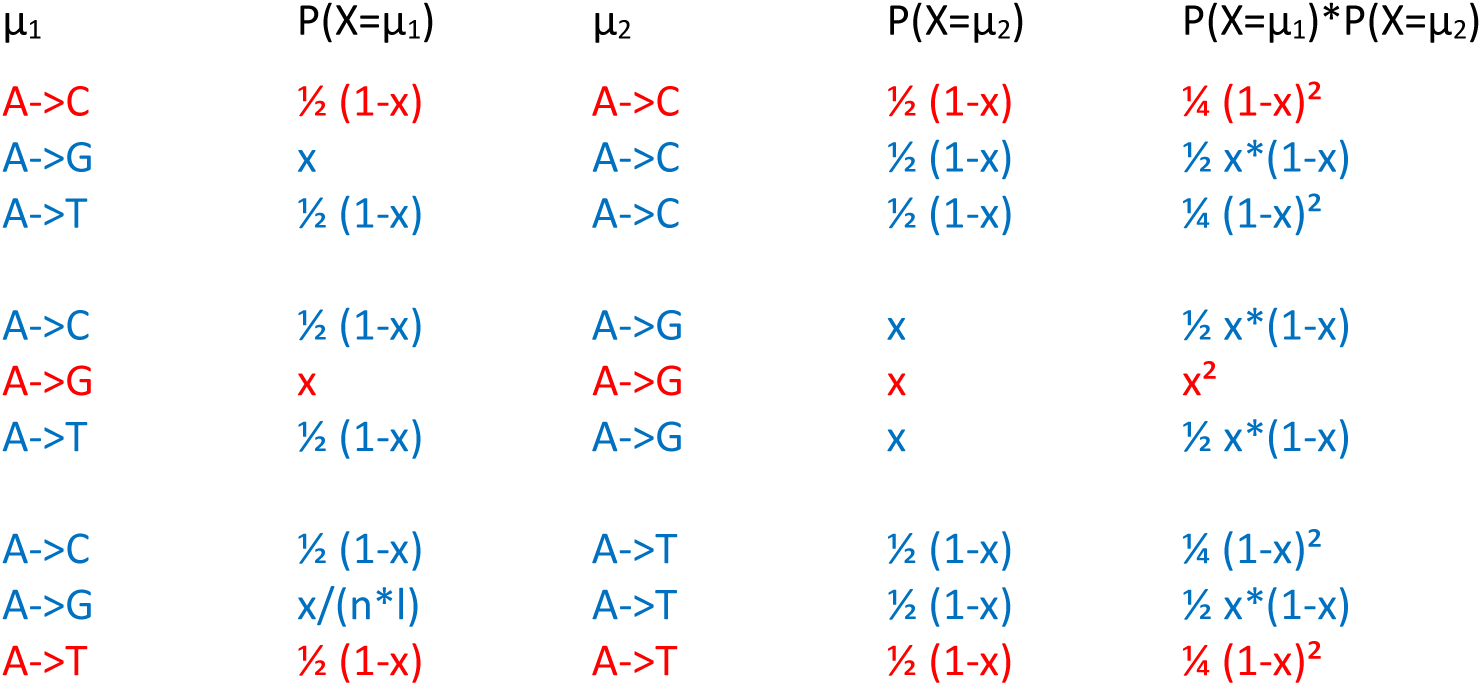

The pattern shown above for possible outcomes starting from ‘A’ is also applicable to the three other nucleotides. The probability that the same point mutation occurs twice at the same site (red cases) is:

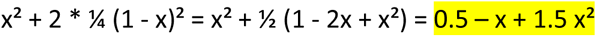

Note that identical mutations occurring independently at the same site in different individuals within a large population go unnoticed.

The probability of different point mutations arising at the same site (blue cases) is:

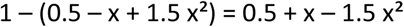

If two distinct point mutations occur at the same site in the same copy in different individuals within a large population, three different nucleotides will segregate at that site.

The first mutation creates a polymorphism at the affected site, and the mutant nucleotide may increase in frequency (f) within the population. Does this alter the probabilities of the outcomes following a second mutation at the same site in the same copy?

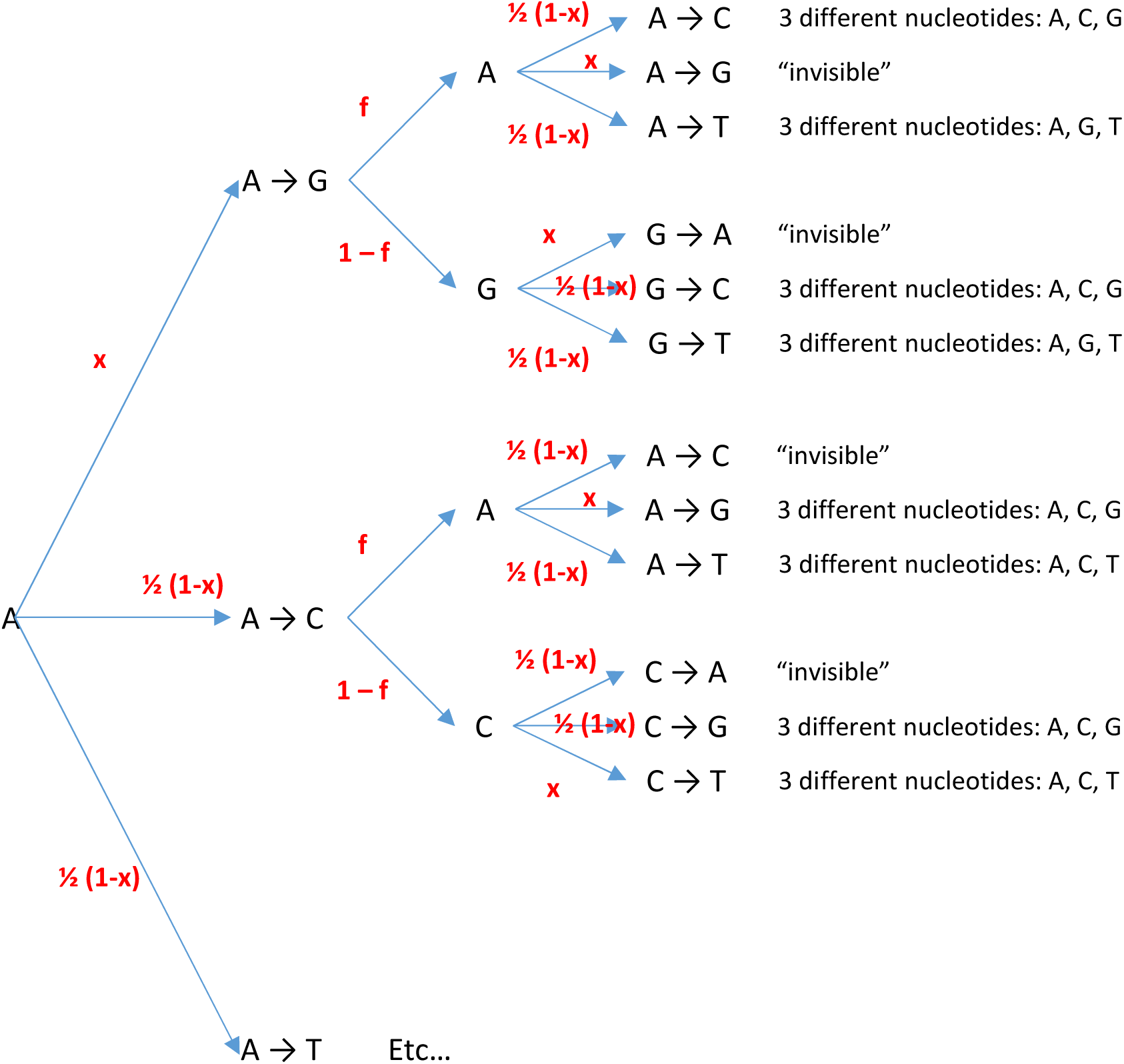

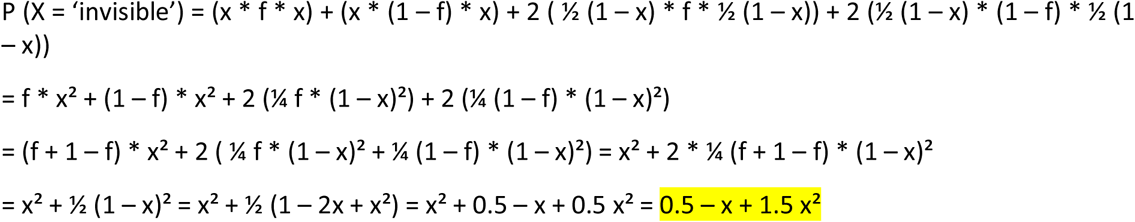

Thus, the probabilities for ‘invisible’ events and those where three different nucleotides segregate at the same site after the second mutation are independent of the mutant allele frequency.

If two mutations hit the same site in the same copy, the probabilities of the outcomes are:

1. **Invisible**: (0.5 – x + 1.5 x²)
2. **Three nucleotides segregating at the same site** in one copy: (0.5 + x – 1.5 x²)

If two mutations occur at the same site in different copies, assuming both copies initially have the same nucleotide, two outcomes are possible:

1. **Shared polymorphism**: Identical mutations result in the same nucleotides segregating at the same site in different copies. Probability: (0.5 – x + 1.5 x²)
2. **Copy-specific polymorphisms**: Different mutations differ result in unique polymorphisms at the same site in each copy. Probability: (0.5 + x – 1.5 x²)

If the two copies initially had different nucleotides due to an ancient mutation, the outcomes depend on whether the differences are transitions or transversions.

a) The copies differ by a transition: A & G or C & T

| 578 | $\mu_1$ | $P(X=\mu_1)$ | $\mu_2$ | $P(X=\mu_2)$ | $P(X=\mu_1)*P(X=\mu_2)$ |
| --- | --- | --- | --- | --- | --- |
| 579 | A->C | $\frac{1}{2} (1-x)$ | G->A | x | $\frac{1}{2} x*(1-x)$ |
| 580 | A->G | x | G->A | x | $x^2$ |
| 581 | A->T | $\frac{1}{2} (1-x)$ | G->A | x | $\frac{1}{2} x*(1-x)$ |
| 582 |  |  |  |  |  |
| 583 | A->C | $\frac{1}{2} (1-x)$ | G->C | $\frac{1}{2} (1-x)$ | $\frac{1}{4} (1-x)^2$ |
| 584 | A->G | x | G->C | $\frac{1}{2} (1-x)$ | $\frac{1}{2} x*(1-x)$ |
| 585 | A->T | $\frac{1}{2} (1-x)$ | G->C | $\frac{1}{2} (1-x)$ | $\frac{1}{4} (1-x)^2$ |
| 586 |  |  |  |  |  |
| 587 | A->C | $\frac{1}{2} (1-x)$ | G->T | $\frac{1}{2} (1-x)$ | $\frac{1}{4} (1-x)^2$ |
| 588 | A->G | x | G->T | $\frac{1}{2} (1-x)$ | $\frac{1}{2} x*(1-x)$ |
| 589 | A->T | $\frac{1}{2} (1-x)$ | G->T | $\frac{1}{2} (1-x)$ | $\frac{1}{4} (1-x)^2$ |

A ‘shared polymorphism’ (red) arises with probability P = x². With probability P = (1 – x²), two copy-specific polymorphisms occur at the same site in both copies.

b) The copies differ by a transversion: A & C, A & T, G & C, or G & T

| 595 | $\mu_1$ | $P(X=\mu_1)$ | $\mu_2$ | $P(X=\mu_2)$ | $P(X=\mu_1)*P(X=\mu_2)$ |
| --- | --- | --- | --- | --- | --- |
| 596 | A->C | $\frac{1}{2} (1-x)$ | C->A | $\frac{1}{2} (1-x)$ | $\frac{1}{4} (1-x)^2$ |
| 597 | A->G | x | C->A | $\frac{1}{2} (1-x)$ | $\frac{1}{2} x*(1-x)$ |
| 598 | A->T | $\frac{1}{2} (1-x)$ | C->A | $\frac{1}{2} (1-x)$ | $\frac{1}{4} (1-x)^2$ |
| 599 |  |  |  |  |  |
| 600 | A->C | $\frac{1}{2} (1-x)$ | C->G | $\frac{1}{2} (1-x)$ | $\frac{1}{4} (1-x)^2$ |
| 601 | A->G | x | C->G | $\frac{1}{2} (1-x)$ | $\frac{1}{2} x*(1-x)$ |
| 602 | A->T | $\frac{1}{2} (1-x)$ | C->G | $\frac{1}{2} (1-x)$ | $\frac{1}{4} (1-x)^2$ |
| 603 |  |  |  |  |  |
| 604 | A->C | $\frac{1}{2} (1-x)$ | C->T | x | $\frac{1}{2} x*(1-x)$ |
| 605 | A->G | x | C->T | x | $x^2$ |
| 606 | A->T | $\frac{1}{2} (1-x)$ | C->T | x | $\frac{1}{2} x*(1-x)$ |

A ’shared polymorphism’ (red) occurs with a probability of P = ¼ (1 – x)². With a probability of P = 1 – (¼ (1 – x)²), we obtain two copy-specific polymorphisms at the same site in both copies.

The relative frequencies of scenarios a) and b) are themselves determined by the probabilities of transitions and transversions. Specifically, in x instances, genes with a transition difference serve as the starting point. Conversely, in (1 – x) instances, genes with a transversion difference are the starting point.

Therefore, the probability of obtaining a shared polymorphism starting with a transition difference is P = x³, and the probability when starting with a transversion difference is P = ¼ (1-x)³. This sums up to a total probability of P = ¼ (1 – 3x + 3x² + 3x³) for copies that initially had different nucleotides at the relevant position. Conversely, the total probability of ending up with two different copy-specific polymorphisms at the same position in both copies is P = ¾ (1 + x - x² - x³).

The exact number of mutable sites in our dataset is uncertain. An initial approximation can be based on the number of synonymous sites, though many synonymous sites in *IGMT*s remain unmutated. Conversely, some non-synonymous sites exhibit polymorphisms or substitutions.

To address this, we have explored scenarios ranging from zero to 100% identical nucleotides, considering that the events of ‘two mutations of identical nucleotides at the same position’ and ‘two mutations of distinct nucleotides at the same position’ are mutually exclusive.

Probabilities of different outcomes for two mutations occurring at the same site X (Overview):

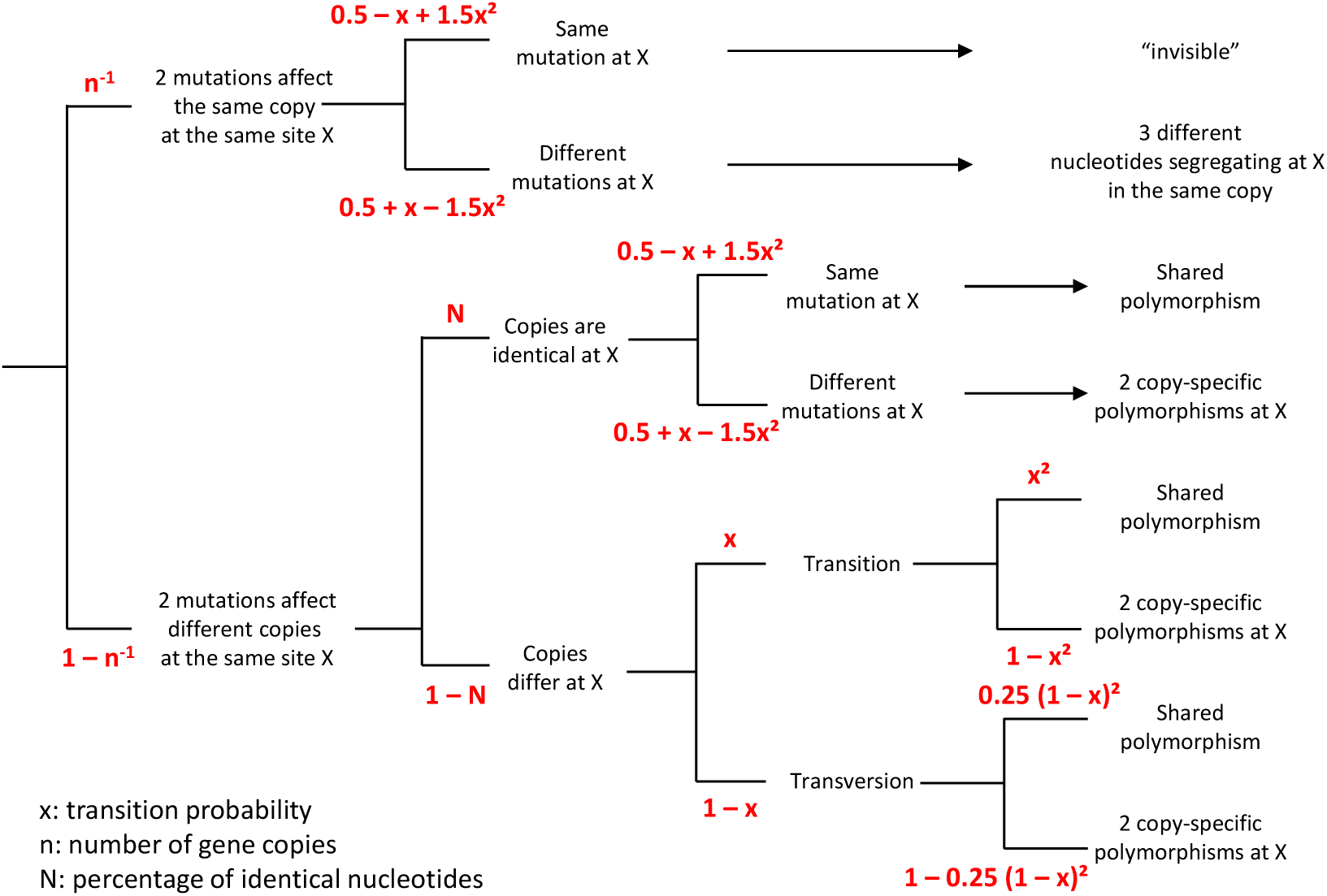

## Supplementary Figures and Tables

**Fig. S1.**
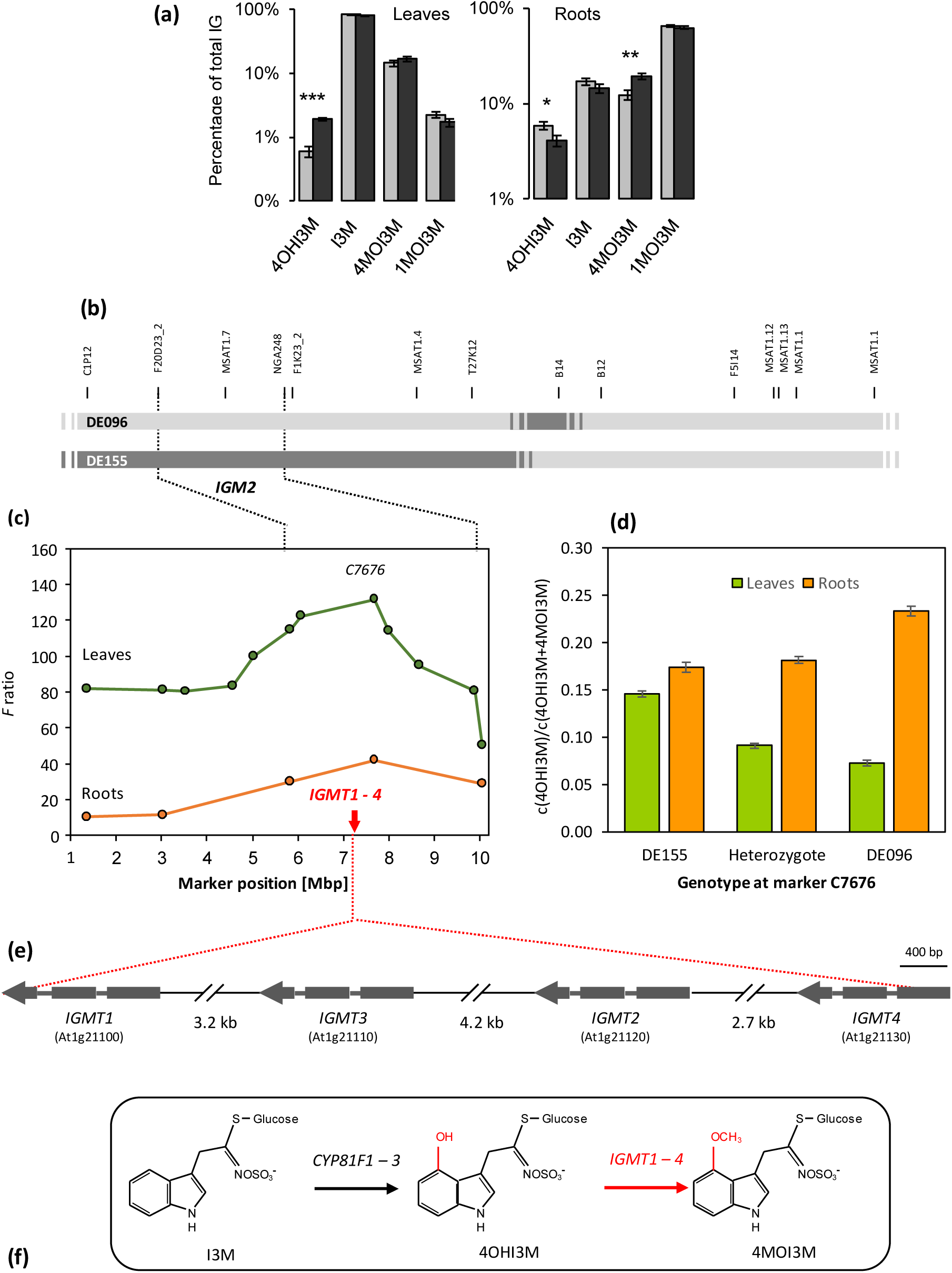
Fine mapping of *Indole Glucosinolate Modifier 2*. a) Indole glucosinolate phenotypes of near-isogenic lines DE096 (light gray) and DE155, shown as percentages of total indole glucosinolate. b) Genotypes of DE096 and DE155 on chromosome 1. DE096 carries parental alleles from the Arabidopsis accession Ei-2 (light gray) in the *IGM2* candidate (dashed lines) region, whereas DE155 carries alleles from Da(1)-12 (dark gray). DE096 and DE155 marker genotypes do not differ on chromosomes 2 – 5. c) Fine mapping. We fine-mapped the leaf QTL as the arcsine square root-transformed proportion of 4OHI3M among modified indole glucosinolates [c4OHI3M x c(4OHI3M + 4MOI3M)^-1^] in 184 F_2_ progeny of a cross between near isogenic lines DE096 and DE155 with twelve markers. Marker positions refer to their physical position in the Col-0 reference genome. Statistical support (green line) for the presence of a QTL peaked close to *IGMT1-4*. Subsequently, we confirmed the presence of a similar QTL in roots (orange line) with a subset of markers in an independent experiment with 184 F_2_ progeny. d) Estimated proportion (± SEM) of 4OHI3M by genotype at marker C7676 in leaves (green) and roots (orange). e) Organization of IGMT1 – 4. Exon-intron structures are based on the Col-0 accession and drawn to scale, with the intervening sequence shortened. The gene order is the same in all investigated accessions. f) Chemical structures of indole glucosinolates and reactions catalyzed by CYP81F1 – 3 and IGMT 1 – 4. *CYP81F1 – 3* gene products hydroxylate indol-3ylmethyl glucosinolate (I3M) at the 4-position of the indole ring, producing 4-hydroxy-I3M (4OHI3M). This compound then serves as the substrate for *IGMT1 – 4* gene products, which generate 4-methoxy-I3M (4MOI3M).

**Fig. S2.**
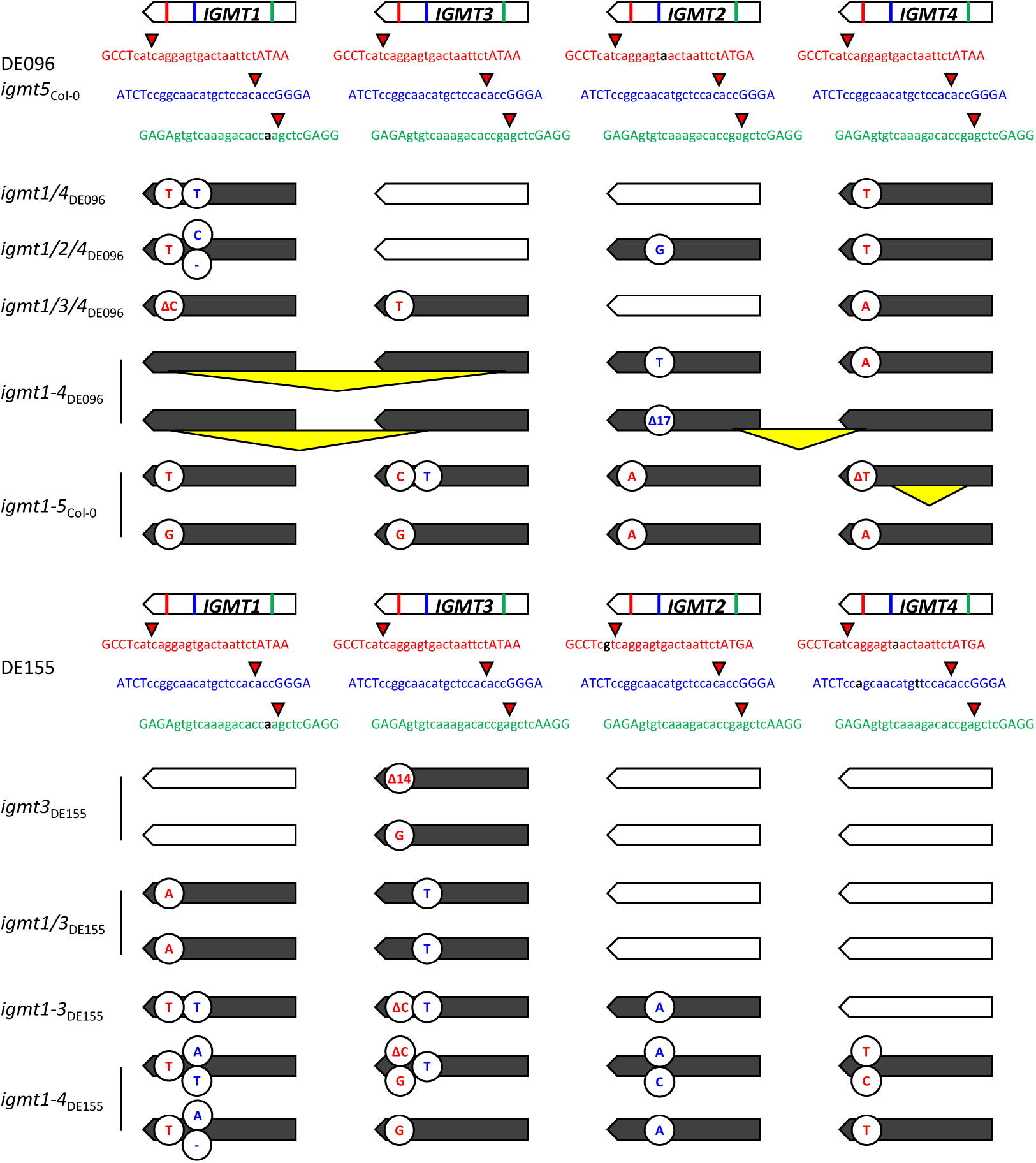
Schematic representation of *IGMT* wild types and mutants. Locations of sgRNAs (red, blue and green colors) in *IGMT1 – 4*, corresponding sequences and flanking nucleotides (capital letters) in DE096, *igmt5*_Col-0_, and DE155. Sequence deviations are shown in black. Mutations rendering genes non-functional are shown in colors corresponding to the different target sites. Insertions are denoted by letters, deletions by a Δ proceeding letters (in case of single-nucleotide deletions) or numbers (in case of multiple nucleotide deletions). Larger deletions, some resulting in fusions between parts of adjacent gene copies, are marked by yellow triangles. Note that some of the target sites were still segregating for polymorphisms when analyzed, leading to heterozygous configurations; however, additional mutations at other target sites ensured non-functionality of the gene products.

**Fig. S3.**
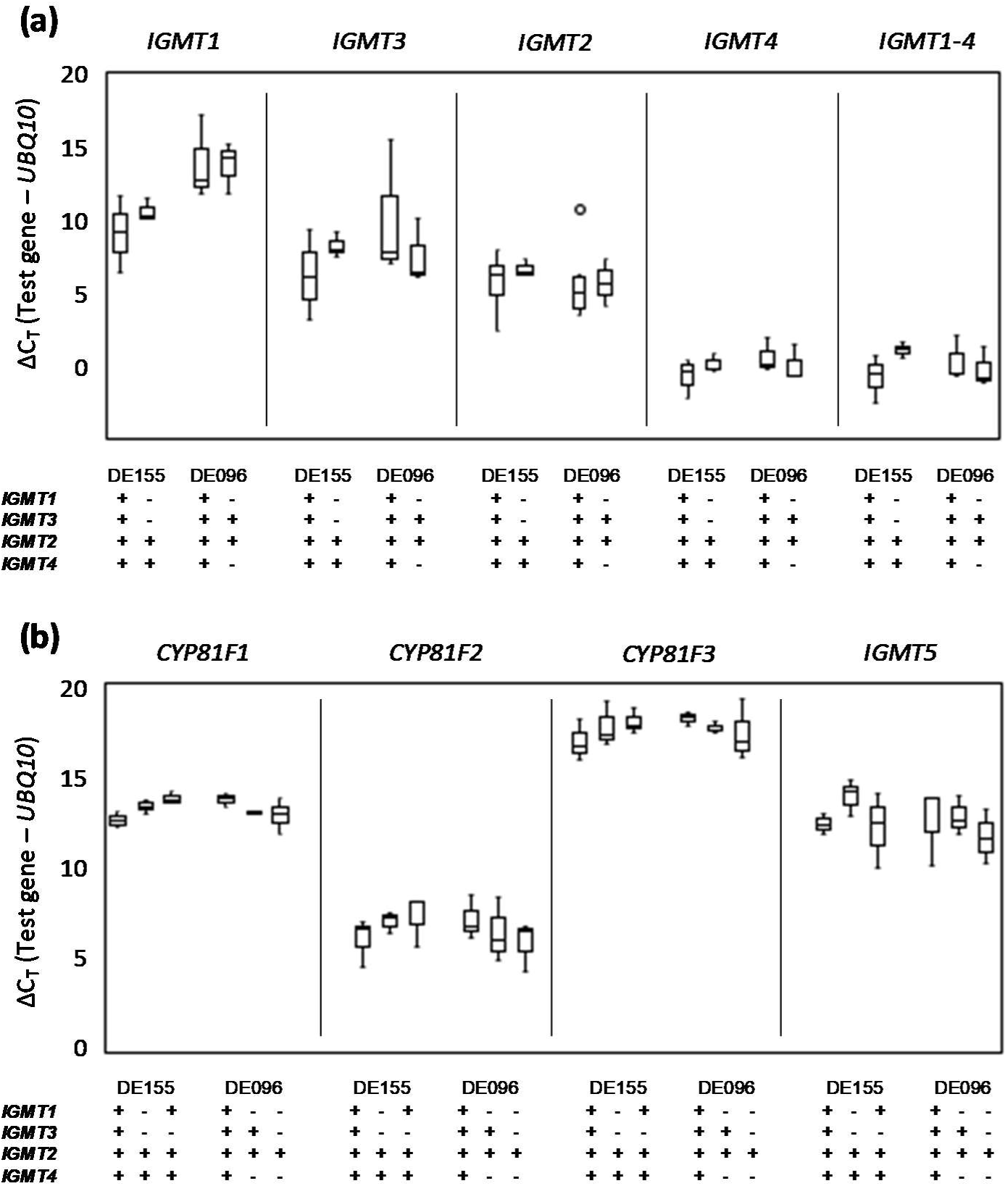
*IGMT* and *CYP81F* transcript levels in wild type and mutant leaves. a) Boxplots showing *IGMT1 – 4* transcript levels in NILs and selected mutant leaves. b) *CYP81F1 – F3* and *IGMT5* transcript levels. Data are presented as median and range of ΔC_T_ values in comparison to the control gene *UBQ10*. Genotypes at *IGMT1 – 4* are indicated, with “-“ for defective and “+” for functional genes.

**Fig. S4.**
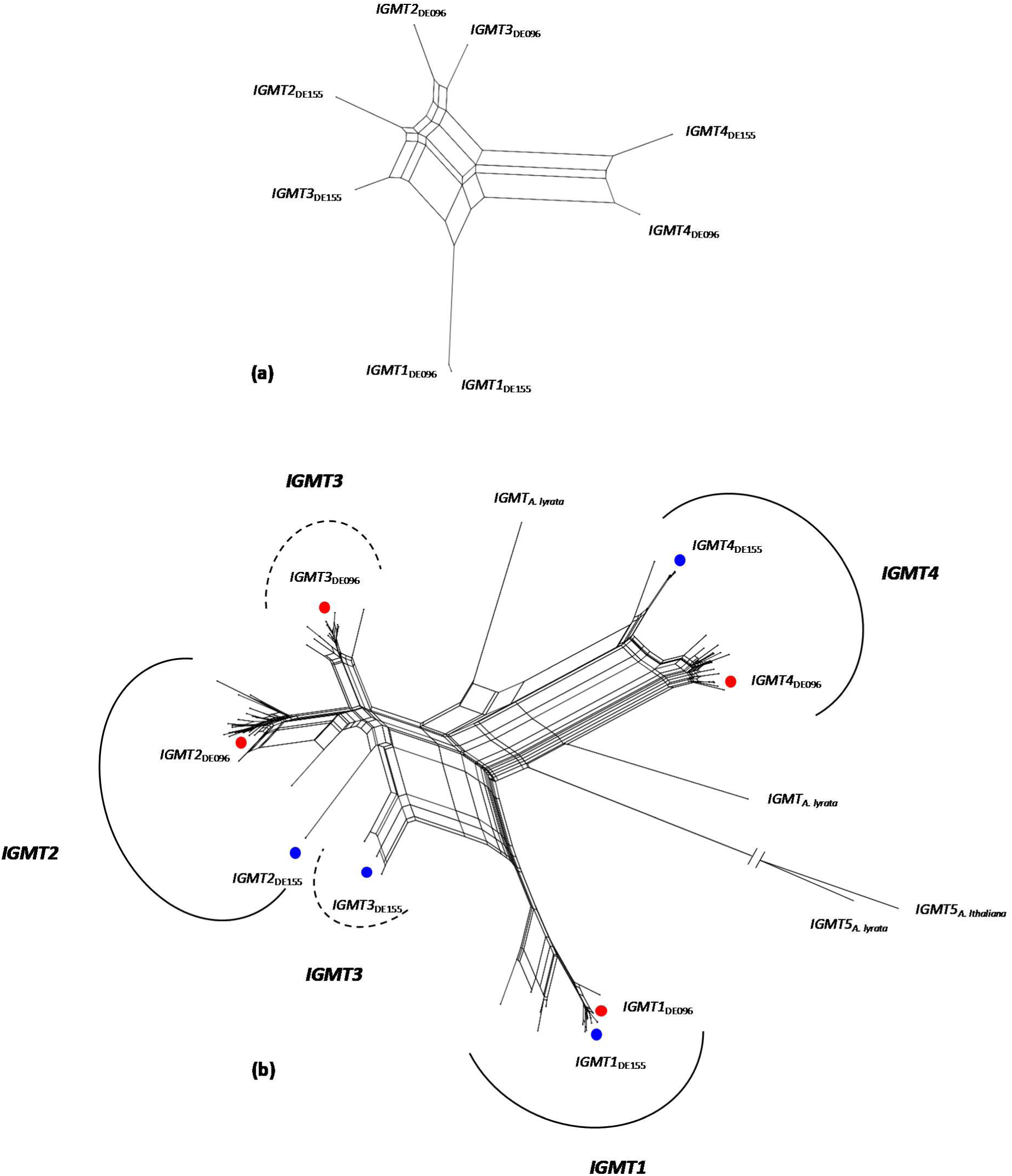
Genealogical networks of *IGMT* genes. a) Network based on DE155 and DE096 *IGMT* genes. b) Network based on *IGMT1 – 4* genes from 28 Arabidopsis accessions, IGMT5, and corresponding sequences from *A. lyrata*. Colored dots indicate the positions of DE155 (blue) and DE096 (red). The *IGMT5* branch was shortened to 1/5 of its original length. Note that there are two clusters with *IGMT3* genes.

**Fig. S5.**
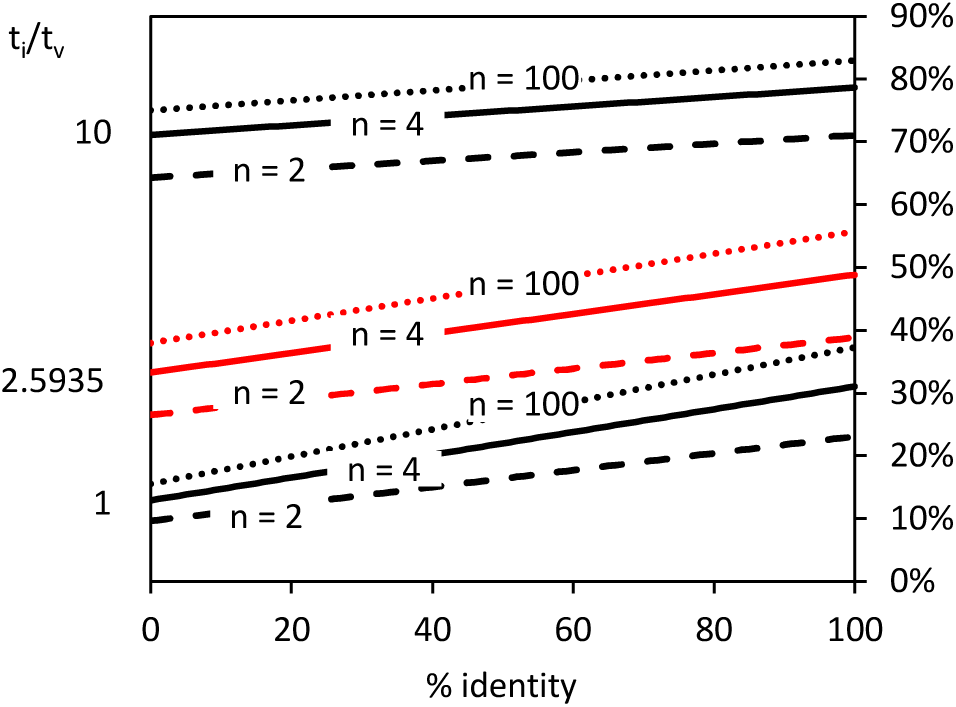
Expected proportion of shared among visible types of polymorphisms for different transition/transversion (t_i_/t_v_) ratios and copy numbers (n). % identity refers to cases where two mutations hit different copies at the same site, which may carry either identical or different nucleotides. Data for Arabidopsis *IGMT1 – 4* are indicated by a solid red line.

**Fig. S6.**
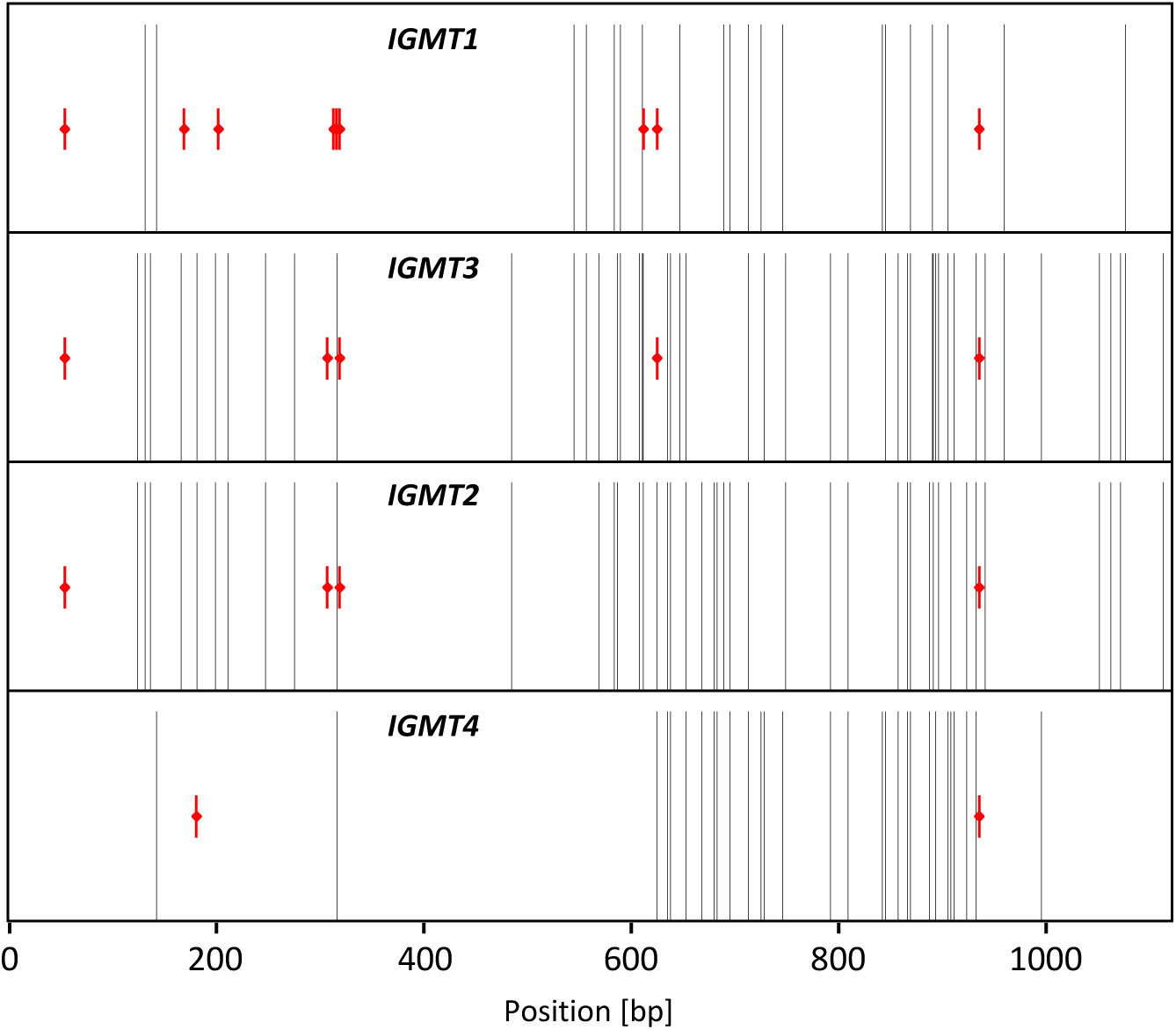
Position of derived amino acids and shared polymorphisms in *IGMT1 – 4* coding sequences. A polymorphism is considered ‘shared’ when the same nucleotides segregate at the same site in two or more gene copies. Intervals flanked by two adjacent shared polymorphisms (black) are significantly larger when they contain derived amino acids (red). Positions are shown in base pairs.

**Fig. S7.**
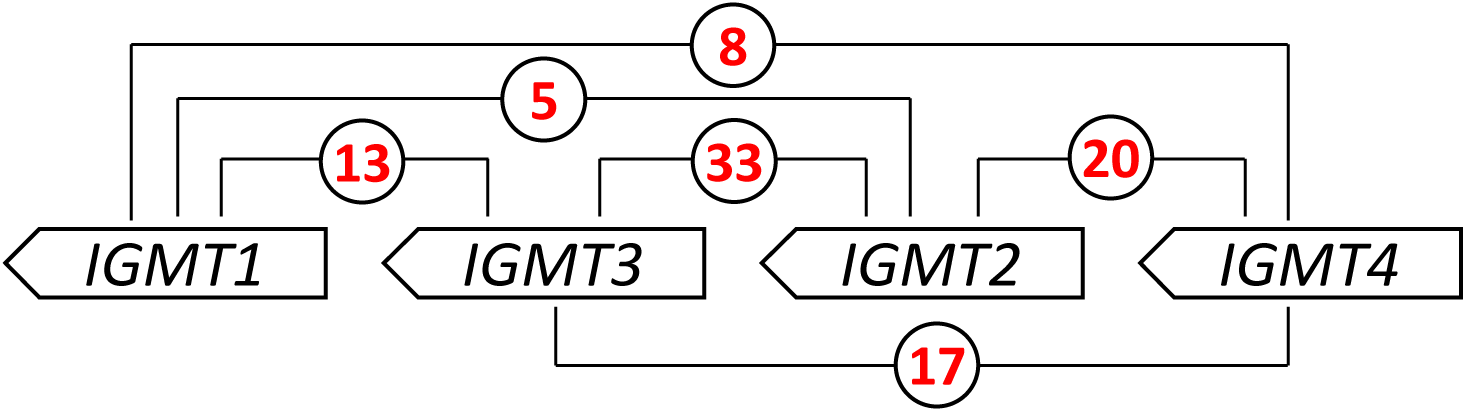
Number of shared polymorphisms in pairwise comparisons of *IGMT* genes. Note that shared polymorphisms are more frequent when genes are in close proximity. Also, the two central genes exhibit the highest number of shared polymorphisms.

**Table S1.**
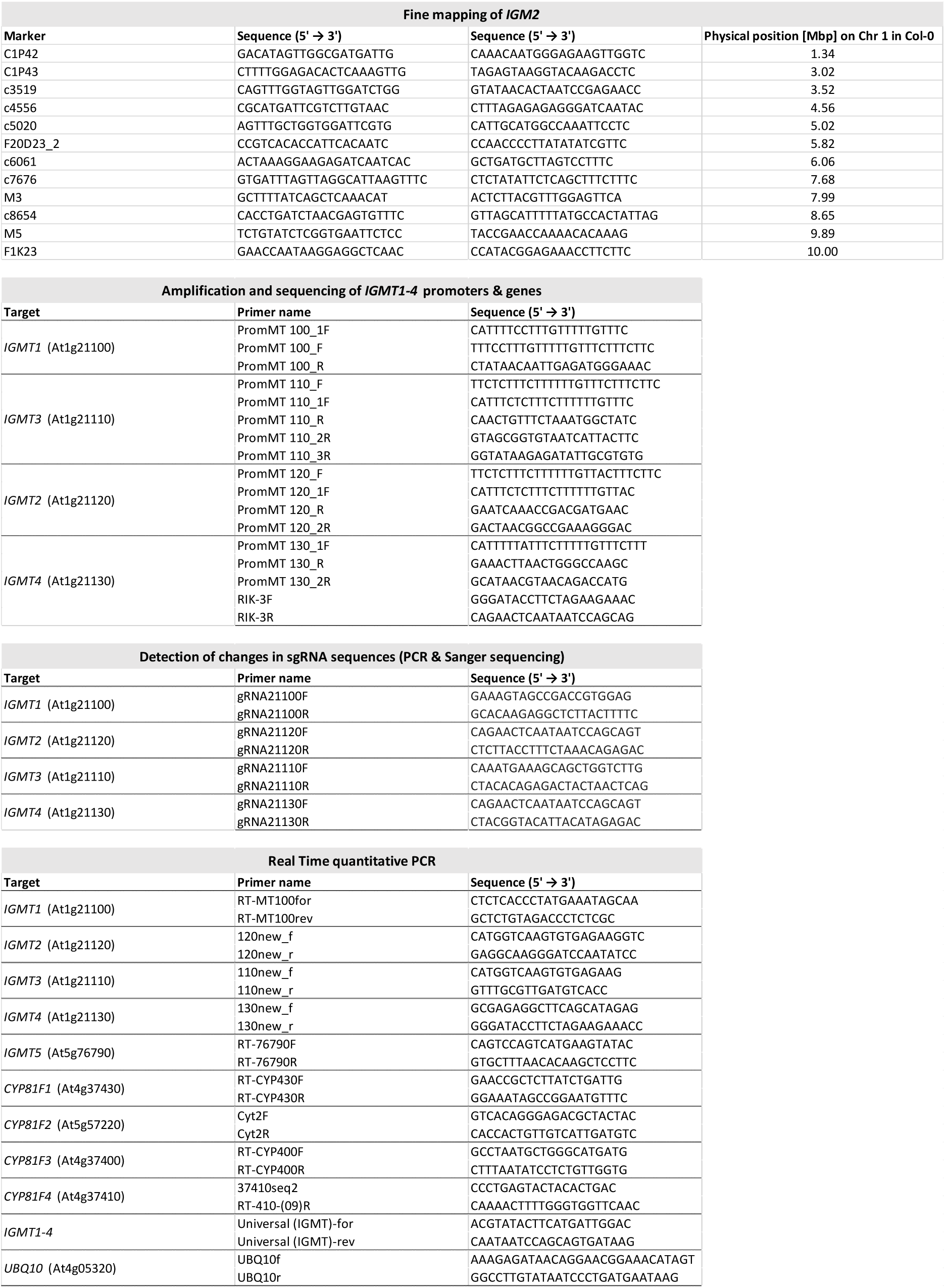
Primer sequences used in this study.

**Table S2.**
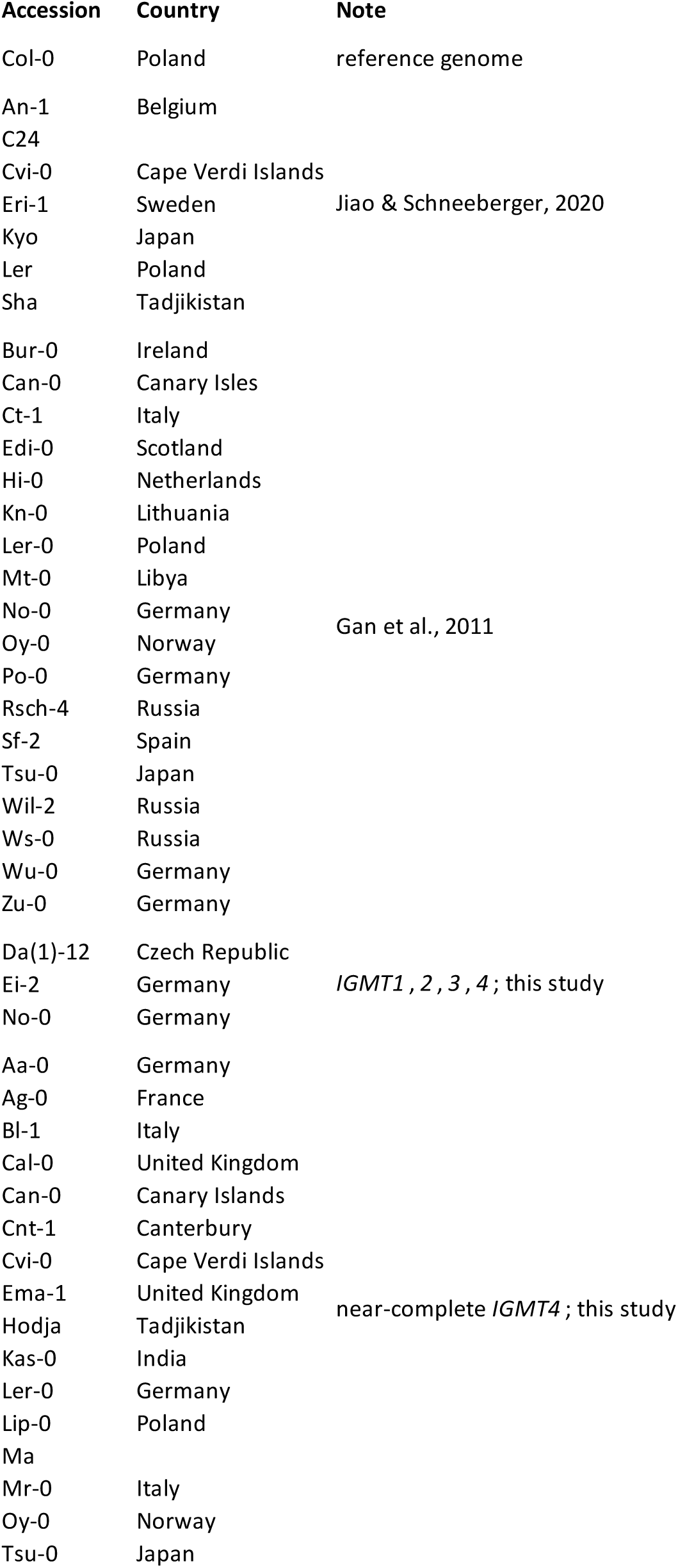
Arabidopsis accessions used in this study.

**Table S3.**
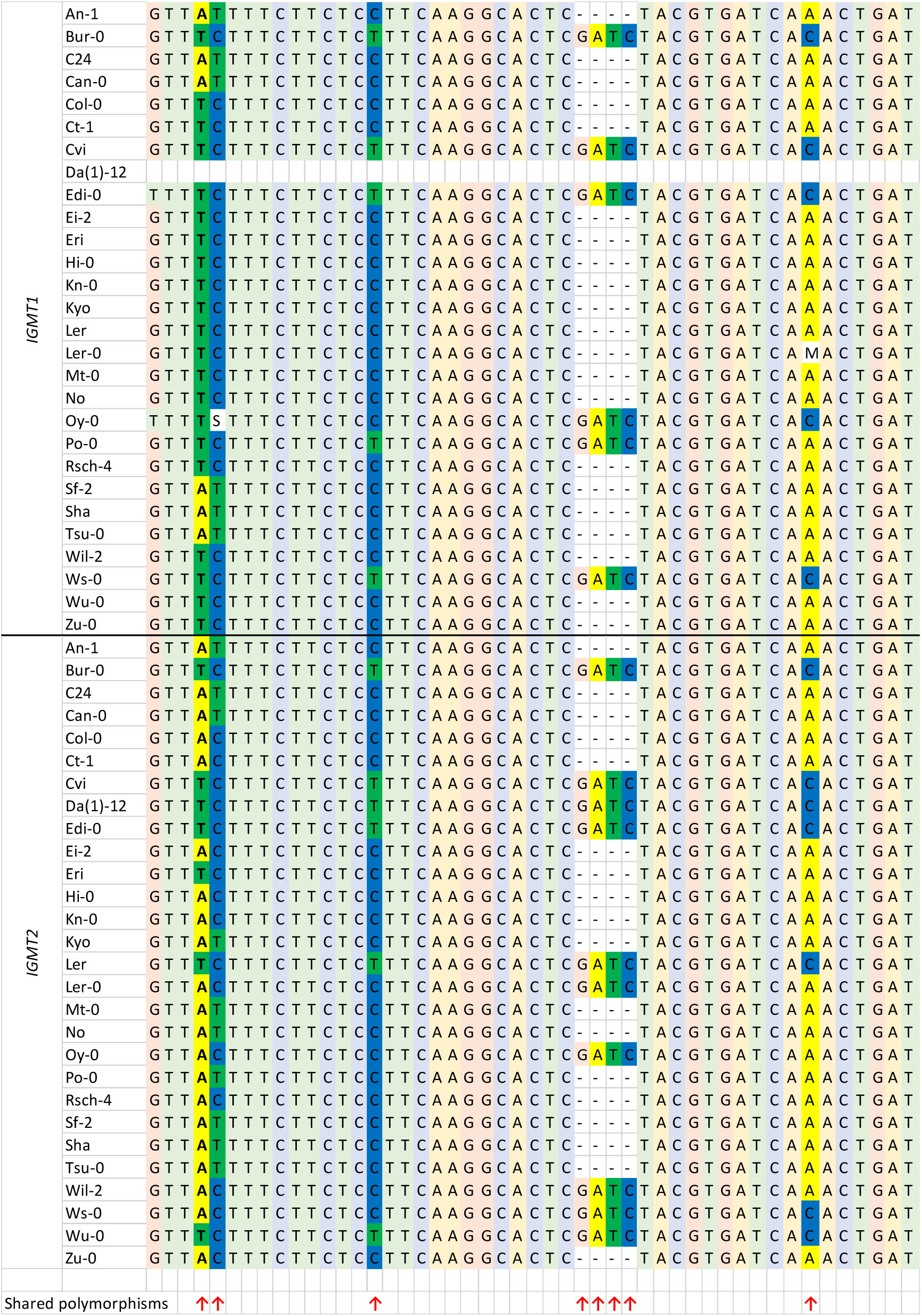
Shared polymorphisms in the 5’UTR of *IGMT1* and *2*.

**Table S4.**
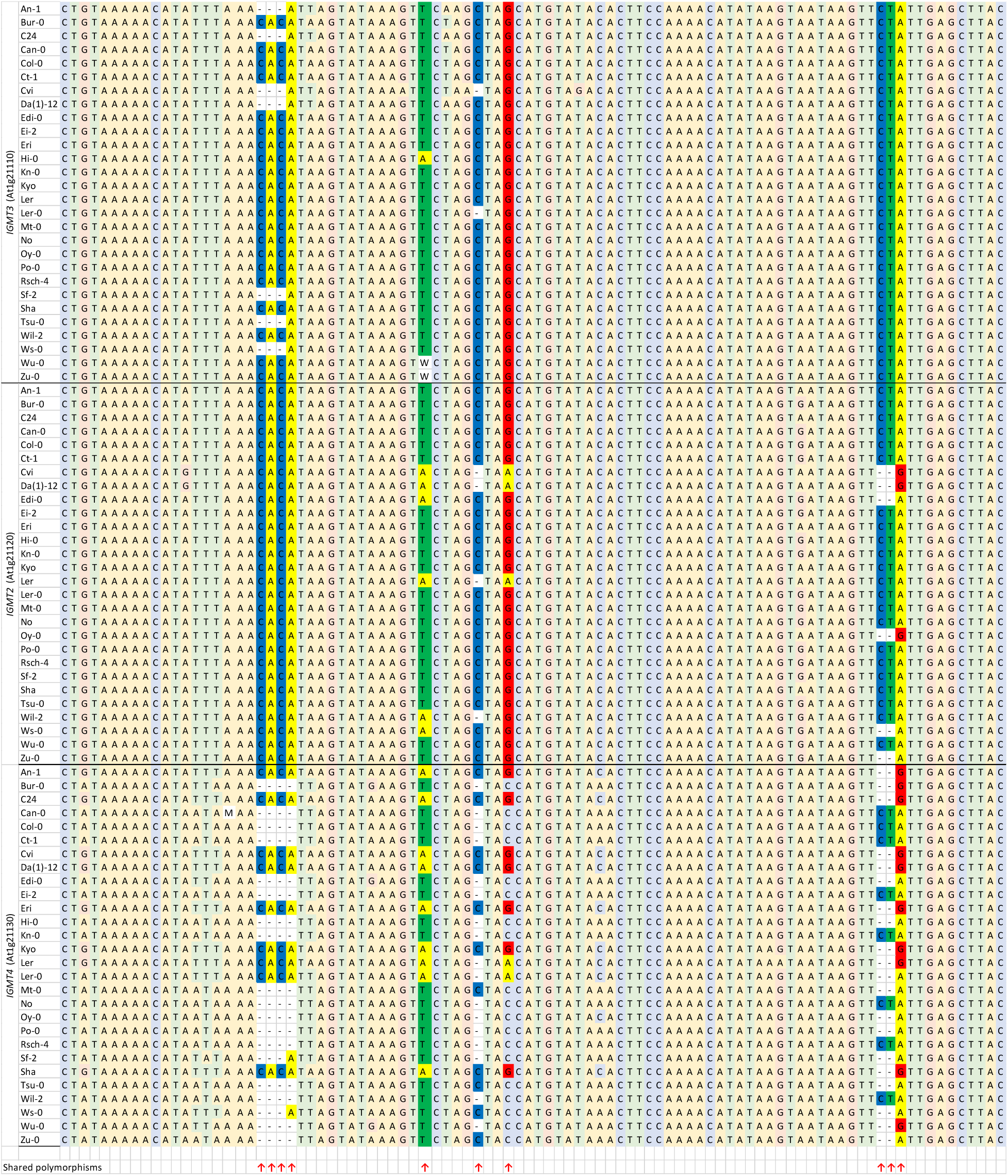
Shared polymorphisms in an *IGMT* intron.

## Notes

### Competing Interest Statement

The authors have declared no competing interest.

### Summary of Updates

We have revised the abstract and large parts of the introduction and discussion, added results suggesting balancing selection on IGMT4, and updated figures and references accordingly.

